# Hepatocyte Dedifferentiation Profiling In Alcohol-Related Liver Disease Identifies CXCR4 As A Driver Of Cell Reprogramming

**DOI:** 10.1101/2023.04.04.535566

**Authors:** Beatriz Aguilar-Bravo, Silvia Ariño, Delia Blaya, Elisa Pose, Raquel A. Martinez García de la Torre, María U Latasa, Celia Martínez-Sánchez, Laura Zanatto, Laura Sererols-Viñas, Paula Cantallops, Silvia Affo, Mar Coll, Xavier Thillen, Laurent Dubuquoy, Matías A Avila, Josep Maria Argemi, Arantza Lamas Paz, Yulia A. Nevzorova, Javier Cubero, Ramon Bataller, Juan José Lozano, Pere Ginès, Philippe Mathurin, Pau Sancho-Bru

## Abstract

**Background and Aims:** Loss of hepatocyte identity is associated with impaired liver function in alcohol-related hepatitis (AH). In this context, hepatocyte dedifferentiation gives rise to cells with a hepatobiliary (HB) phenotype expressing biliary and hepatocytes markers and showing immature features. However, the mechanisms and the impact of hepatocyte dedifferentiation in liver disease are poorly understood.

**Methods:** HB cells and ductular reaction (DR) cells were quantified and microdissected from liver biopsies from patients with alcohol-related liver disease (ALD). Hepatocyte-specific overexpression or deletion of CXCR4, and CXCR4 pharmacological inhibition were assessed in mouse liver injury. Patient-derived and mouse organoids were generated to assess plasticity.

**Results:** Here we show that HB and DR cells are increased in patients with decompensated cirrhosis and AH, but only HB cells correlate with poor liver function and patients’ outcome. Transcriptomic profiling of HB cells revealed the expression of biliary-specific genes and a mild reduction of hepatocyte metabolism. Functional analysis identified pathways involved in hepatocyte reprogramming, inflammation, stemness and cancer gene programs. CXCR4 pathway was highly enriched in HB cells, and correlated with disease severity and hepatocyte dedifferentiation. *In vitro*, CXCR4 was associated with biliary phenotype and loss of hepatocyte features. Liver overexpression of CXCR4 in chronic liver injury decreased hepatocyte specific gene expression profile and promoted liver injury. CXCR4 deletion or its pharmacological inhibition ameliorated hepatocyte dedifferentiation and reduced DR and fibrosis progression.

**Conclusions:** This study shows the association of hepatocyte dedifferentiation with disease progression and poor outcome in AH. Moreover, the transcriptomic profiling of HB cells revealed CXCR4 as a new driver of hepatocyte-to-biliary reprogramming and as a potential therapeutic target to halt hepatocyte dedifferentiation in AH.

**Lay summary:** Here we describe that hepatocyte dedifferentiation is associated with disease severity and a reduced synthetic capacity of the liver. Moreover, we identify the CXCR4 pathway as a driver of hepatocyte dedifferentiation and as a therapeutic target in alcohol-related hepatitis.

## INTRODUCTION

Alcohol-related hepatitis (AH) is an acute-on-chronic liver injury condition developed in patients with underlying alcohol-related liver disease (ArLD) and recent alcohol intake. AH is characterized by cholestasis, cell damage, inflammatory cell infiltration, hepatocellular failure and loss of hepatocyte identity [1–3]. Moreover, patients with AH are characterized by the expansion of the ductular reaction (DR), which is associated with liver failure and high short-term mortality [1,4].

Both hepatocytes and biliary cells present a high plasticity potential, which is essential for their response to stress and injury. In liver injury, hepatocyte plasticity plays a key role in cell response to stress and is critical for cell proliferation and liver regeneration [5–7]. In this context, the biliary epithelium gives rise to immature cells and the expansion of DR, which emerges as a regenerative response of the liver to sustain the biliary compartment [8,9]. However, in the context of advanced liver disease, DR expansion and hepatocyte reprogramming become a maladaptive regenerative response of the liver and are associated with disease progression [10,11].

Hepatocyte-to-biliary reprogramming involves a progressive acquisition of biliary features by hepatocytes while reducing hepatocyte characteristics and functional competence, giving rise to hepatobiliary cells HB. Several histological studies have shown the presence of HB expressing both, hepatocyte and biliary markers in cirrhosis, acute liver failure (ALF) and cholestatic liver diseases [10,12]. In AH, we have shown that the transforming growth factor beta (TGFβ) pathway plays a key role promoting hepatocyte reprogramming [2]. In the same line, the activation of the Hippo/YAP pathway is involved in adult hepatocyte loss of identity in patients with ALF and AH [13–15]. In mice, lineage tracing studies have provided evidence for hepatocyte plasticity in chronic injury models [16,17]. In this context, Notch, YAP and Wnt pathways promote hepatocyte dedifferentiation and hepatocyte-to-biliary transdifferentiation [16,18][7,9,16]. In addition, hepatocytes can be the source of both intrahepatic cholangiocarcinoma [19], thus suggesting the association of hepatocyte-to-biliary reprogramming with tumorigenesis. While hepatocyte reprogramming has been extensively studied in animal models, its impact in human liver disease and the molecular mechanisms driving this event are not well understood.

These studies revealed the importance of preserving tight control of hepatocyte reprogramming in response to injury to maintain liver function and tissue repair, and underline cell reprogramming as a potential therapeutic target in chronic liver disease. In this study, we aimed to dissect the molecular mechanisms regulating hepatocyte-to-biliary reprogramming inducing the appearance of HB cells in ArLD. We reveal that HB cells constitute a hallmark of AH, correlating with disease severity and reduced synthetic capacity of the liver. Moreover, we describe the transcriptomic and functional profile of human HB cells identifying CXCR4 as a signaling pathway underlying hepatocyte-to-biliary reprogramming and as a potential therapeutic target to revert hepatocyte dedifferentiation in AH.

## METHODS

### Patient information

Liver biopsies from patients with ArLD admitted to the Liver Unit of the Hospital Clinic of Barcelona were used. Signed informed consent was obtained from all the patients, and the study was approved by the Ethics Committee of the Hospital Clinic. The ArLD cohort included patients in different disease stages: pre-cirrhosis (n=3), compensated cirrhosis (n=4), decompensated cirrhosis (n=13) and AH patients (n=15). Associated clinical and biochemical parameters are shown in Table 1. Laser capture microdissection (LCM) was performed in explants of patients with AH admitted to the Hôpital Claude Huriez of Lille (France) who underwent liver transplantation. Patient information was previously described [20]. Informed consent was obtained from all the patients (QuickTrans (NCT01756794) and TargetOH (CPP 14/67) studies).

## RESULTS

### Hepatobiliary cells are a hallmark of alcohol-related hepatitis and correlate with poor outcomes in these patients

The expansion of DR in patients with AH and its correlation with disease severity has been previously described [10,21]. Moreover, human histological studies have shown the presence of HB cells expressing biliary markers while preserving hepatocyte morphology [10,22]. Here, we envisioned studying HB and DR cells independently to evaluate their association with disease progression. Paraffin-embedded liver biopsies from patients with ArLD were stained for KRT7 (Fig. 1A, Table 1). Both HB and DR cells were increased in patients with decompensated cirrhosis and AH patients as compared with patients with pre-cirrhosis and compensated cirrhosis (Fig. 1B). Interestingly, while the extent of DR was the same in patients with decompensated cirrhosis and AH, the amount of HB cells was increased in patients with AH compared with patients with decompensated cirrhosis (Fig. 1B). Moreover, the amount of HB cells and DR cells correlated positively (r=0.41; p=0.02) (Fig. 1C).

**Fig. 1.**
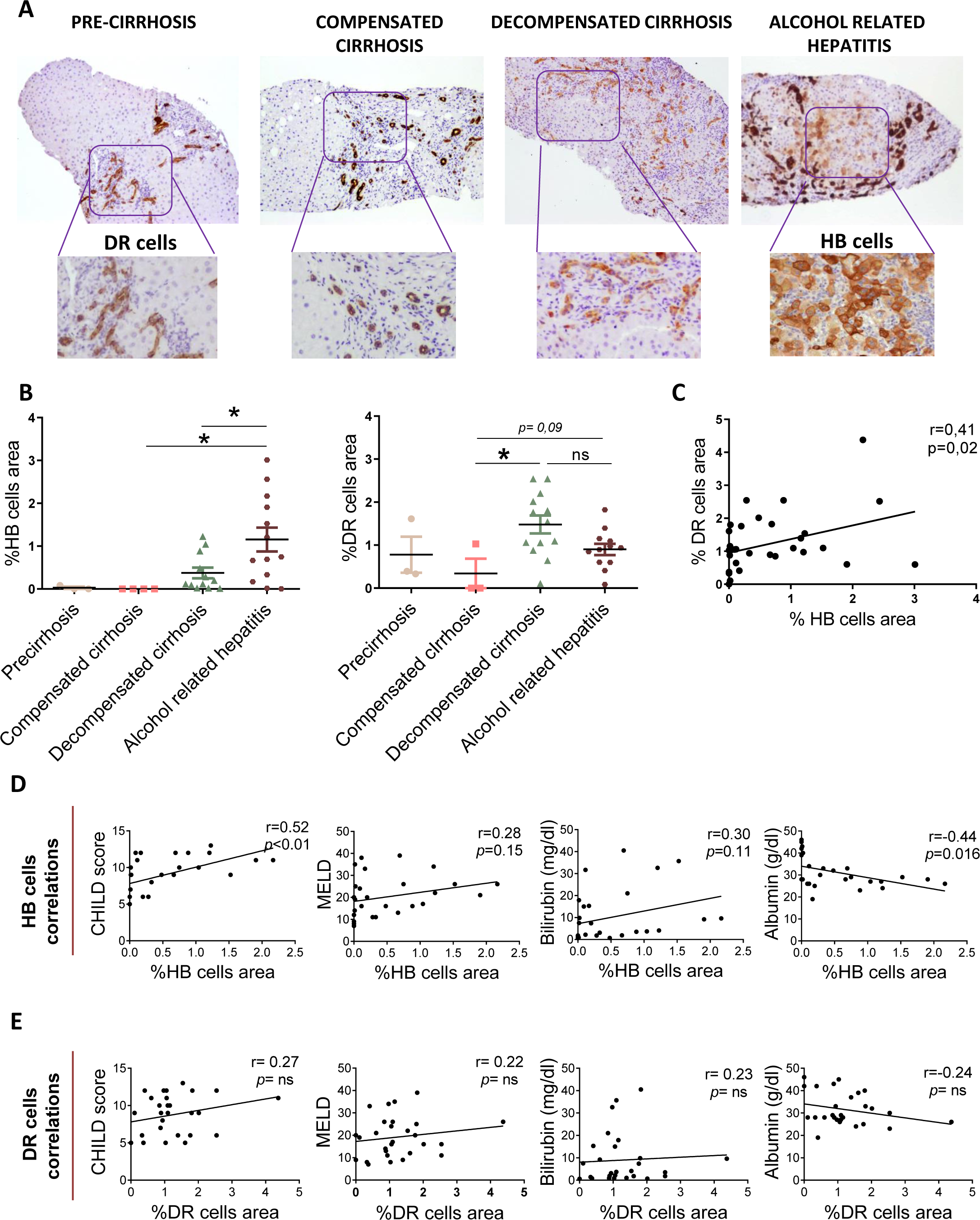
Histological analysis of HB cells and DR cells in the progression of ArLD disease. **(A)** KRT7 staining of liver biopsies from 3 patients with pre-cirrhosis, 4 patients with compensated cirrhosis, 13 patients with decompensated cirrhosis and 15 patients with AH **(B)** Representation of HB cell area (KRT7^+^ hepatocytes) and DR cell area (KRT7^+^ DR cells) along ArLD progression. Significant differences are indicated as \**p*<0.05 (T-student test). **(C)** Correlation between the area of HB cells and DR cells from ArLD biopsies. **(D)** Correlation between HB cells (%area) and **(E)** DR cells (%area) with the clinical and biochemical parameters associated with the ArLD cohort. The regression coefficient (r) and p-value of each correlation are indicated.

Importantly, while there was no significant correlation of DR cells with disease severity, the number of HB cells showed a positive correlation with CHILD PUGH (r=0.52; p<0.01) score and a negative correlation with albumin values (r= -0.4; p=0.03) (Fig. 1D and E). Altogether, these results suggest an association of HB cells but not DR cells with poor liver function in patients with advanced ALD.

### Characterization of the transcriptomic profile of hepatobiliary cells

In order to investigate the gene expression profile of HB cells, we performed Laser capture microdissection (LCM) and RNASeq analysis of HB cells (KRT7^+^ hepatocytes), hepatocytes (KRT7^-^ hepatocytes) and DR cells (KRT7^+^ DR cells) from paraffin-embedded liver sections from four patients with chronic ArLD [20] (Fig. 2A). As shown in Fig. 2B, principal component analysis of transcriptomic data revealed a marked distance between the three populations, indicating important changes in the gene expression profile (Table S1 and S2).

**Fig. 2.**
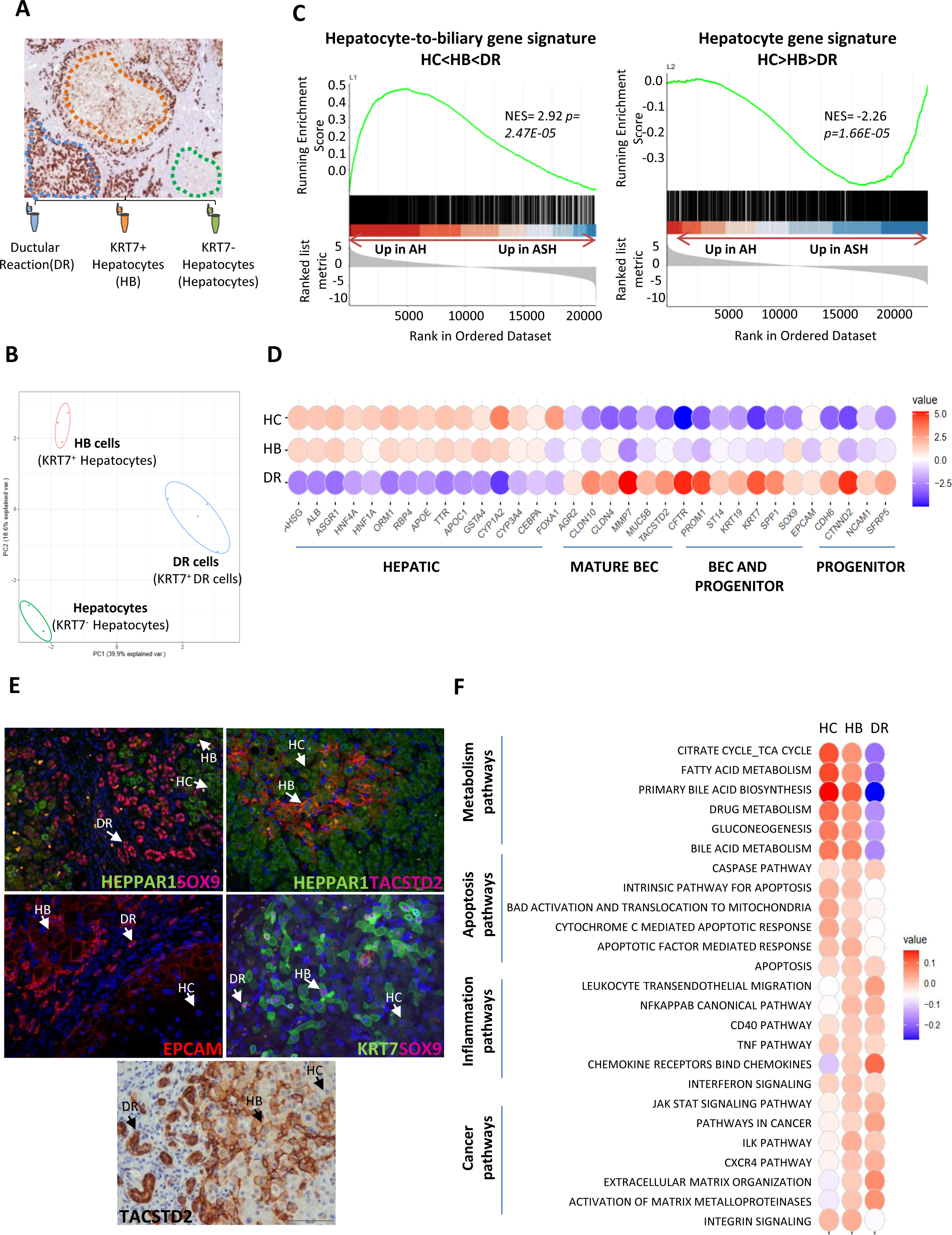
Transcriptomic analysis of microdissected samples from patients with AH. **(A)** KRT7 staining of a liver explant with AH showing the three microdissected populations. DR cells (KRT7^+^ DR cells) in blue (n=4), HB cells (KRT7^+^ hepatocytes) in orange (n=3) and hepatocytes (KRT7^-^ hepatocytes) in green (n=2). **(B)** Principal component analysis (PCA) showing the isolated populations that underwent RNA sequencing. Each sample is positioned in the two-dimensional space according to its RNA expression. **(C)** GSEA of the hepatocyte-to-biliary gene signature (hepatocytes<HB<DR) (upper panel) and the hepatocyte gene signature (hepatocytes>HB>DR) (bottom panel) in a data set of transcriptomic data from AH vs. ASH patients. Normalized enrichment score (NES) and significance is shown. **(D)** Balloon plot summarizing scaled average expression values of hepatic and biliary genes in the three populations, hepatocytes (HC), HB cells and DR cells. Red and blue dots represent positive and negative enrichment, respectively. **(E)** Representative staining of biliary (EPCAM, KRT7, SOX9 and TROP2) and hepatocyte (HEP-PAR1) markers in paraffin embedded liver sections from patients with AH. Hepatocytes (HC), HB and DR cells are indicated. **(F)** Balloon plot showing selected GSEA in the three populations (HC, HB, DR). Red spots represent activated gene sets and blue spots represent down-regulated gene sets.

Next, we wanted to determine if the expression of biliary genes in hepatocytes was associated with disease severity. Thus we performed a gene set enrichment analysis (GSEA) to evaluate the expression of: 1) Hepatocyte-to-biliary gene signature (genes up-regulated in DR vs. HB vs. Hepatocytes) and 2) Hepatocyte gene signature (genes up-regulated in hepatocytes vs. HB vs. DR) in patients with AH compared to patients with alcohol-related steatohepatitis (ASH) [2] (Fig. 2C, Table S3 and S4). GSEA showed a positive enrichment of the hepatocyte-to-biliary gene signature in patients with AH (NES: 2.92; *p: 2.47E-05*). On the contrary, the hepatocyte gene signature was enriched in patients with ASH (NES: -2.26; *p: 1.66E-05*) (Fig. 2C). These results indicate that patients with AH present an up-regulation of genes associated with hepatocyte-to-biliary reprogramming and loss of hepatocyte identity.

To further analyze the transcriptomic profile of the three microdissected cell populations, well-known hepatocyte, biliary and progenitor key markers were examined. As shown in Fig. 2D and Table S5, there was a mild reduction of hepatocyte-specific genes in HB cells compared to hepatocytes. However, HB cells showed increased expression of biliary and progenitor genes, such as *AGR2*, *CLDN4*, *TACSTD2*, *KRT19* and *SOX9*, among others (Fig. 2D). The expression of biliary-specific markers (EPCAM, KRT7, SOX9 and TROP2) and the hepatocyte marker HEPAR1 was confirmed by immunofluorescence and immunohistochemistry (Fig. 2E) in liver tissue from patients with AH. These results suggest that HB cells show up-regulation of biliary genes while maintaining to some extent the expression of hepatocyte markers.

### Functional analysis of the transcriptome of hepatobiliary cells

To understand the molecular events underlying hepatocyte plasticity, GSEA was used to identify pathways enriched in each population (Fig. 2F and Table S6). The analysis revealed that both hepatocytes and HB cells were enriched in hepatocyte function and metabolic pathways, such as gluconeogenesis or bile acid metabolism, as well as pathways associated with apoptosis. Interestingly, HB cells were enriched in inflammatory pathways, cancer and extracellular matrix, such as the NFKB; JAK-STAT and CXCR4 pathways as well as activation of matrix metalloproteinases.

Next, we sought to perform a more in-depth analysis independently comparing HB cells with hepatocytes or with DR cells. This analysis identified 438 up-regulated genes and 399 down-regulated genes in HB cells vs. hepatocytes. As shown in the volcano plot (Fig. 3A), HB cells showed down-regulation of the Notch-ligand *JAG2* and genes associated with hepatocyte function such as *FOXA1* and *GGT1* among others (Table S1). Genes up-regulated in HB cells included biliary markers, such as *SPP1*, *TACSTD2* and *CLDN4*, epithelial-mesenchymal transition (EMT)-related genes, such as *ZEB2* and *TGFBR1*, and *CXCR4* pathway-associated genes, such as *CXCR4*, *PAG1*, *ITGA4* and *ITGA8* (Fig. 3A, Table S1). Accordingly, comparison of the HB transcriptomic profile to DR cells showed a reduced expression of biliary markers and enrichment of hepatocyte genes (Fig. 3B, Table S2).

**Fig. 3.**
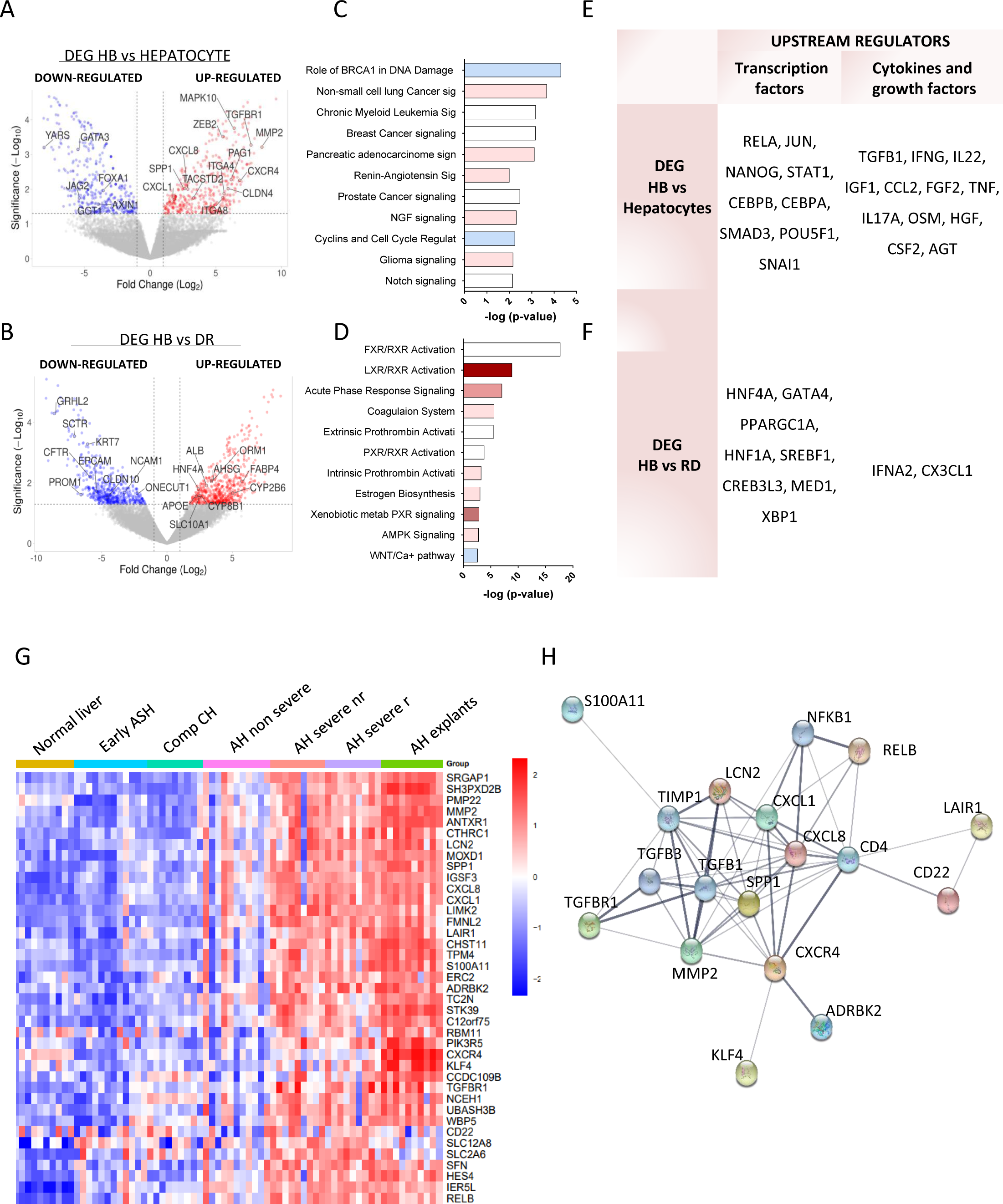
Transcriptomic analysis of HB cells and its association with ArLD severity. **(A)** Volcano plots of differentially expressed genes in HB cells vs. hepatocytes or **(B)** vs. DR cells. Selected genes from each panel are annotated. **(C)** IPA analysis of enriched canonical pathways in HB cells vs. hepatocytes or **(D)** HB cells vs. DR cells. Inhibited pathways are represented in blue, activated pathways are represented in red, and pathways with no status information are represented in white. The pathway significance is shown as -log(p-value). **(E)** IPA prediction of activated upstream regulators in HB cells vs. hepatocytes or **(F)** activated upstream regulators in HB cells vs. DR cells. Upstream regulators with a z-score>2 was annotated. **(G)** Heat map illustrating the HB gene signature along ArLD progression. Red color indicates up-regulated gene expression, while blue color shows decreased gene expression. ASH, alcohol-related steatohepatitis; comp CH, compensated cirrhosis; AH, alcohol-related hepatitis; nr, non-responders; r, responders. Gene correlation with clinical parameters is shown in Table S9 **(H)** Protein interaction network of HB gene signature. 10 out of 39 genes were clustered and 8 additional genes were added by String software. Line thickness indicates the strength of data support.

These two data sets were further analyzed to identify differentially enriched pathways (Fig. 3C, 3D; Table S7 and S8) and upstream regulators (Fig. 3E and 3F). When compared to hepatocytes, HB cells were enriched in pathways related to cancer, inflammation, cell senescence and the CXCL12 signaling pathway (Fig. 3C and Table S7). In addition, upstream regulators predicted to be activated (z-score>2) were identified (Fig. 3E). When compared to DR cells, HB cells were enriched in hepatocyte metabolism pathways and showed a reduction of biliary-associated pathways such as bile secretion, Notch signaling or Wnt-beta catenin signaling (Fig. 3D and Table S8). Upstream regulators were predicted (z-score>2) by IPA (Fig. 3F).

Genes up-regulated in HB cells vs. hepatocytes were evaluated in a cohort of ArLD patients (with whole liver transcriptome data)[2]. As shown in the heatmap (Fig. 3G) and in Table S9, 40 out of 438 genes were up-regulated in severe AH patients compared to non-severe AH and correlated with severity scores. Subsequently, a protein interaction network of these genes revealed the link between CXCR4, the TGFB pathway and CXCL- family members (Fig. 3H). Altogether, these results suggest that expression of inflammation and cancer-associated processes may play a role in hepatocyte reprograming underlying AH pathogenesis.

### *CXCR4* is associated with loss of hepatocyte identity and disease progression

We found that CXCR4-pathway was associated with hepatocyte-to-biliary reprogramming (Fig. 2F) and dramatically up-regulated in HB cells compared to hepatocytes (Fold Change: 112) (Fig. 4A). Moreover, CXCR4 pathway genes were highly expressed in hepatobiliary cells vs. hepatocytes (Table S10). Given the consistent association of CXCR4 expression with the hepatobiliary phenotype and the data reporting the role of CXCR4 in tumor cell plasticity, stemness, inflammation and fibrosis [23,24], we envisioned assessing the role of CXCR4 in hepatocyte-to-biliary reprogramming.

**Fig. 4.**
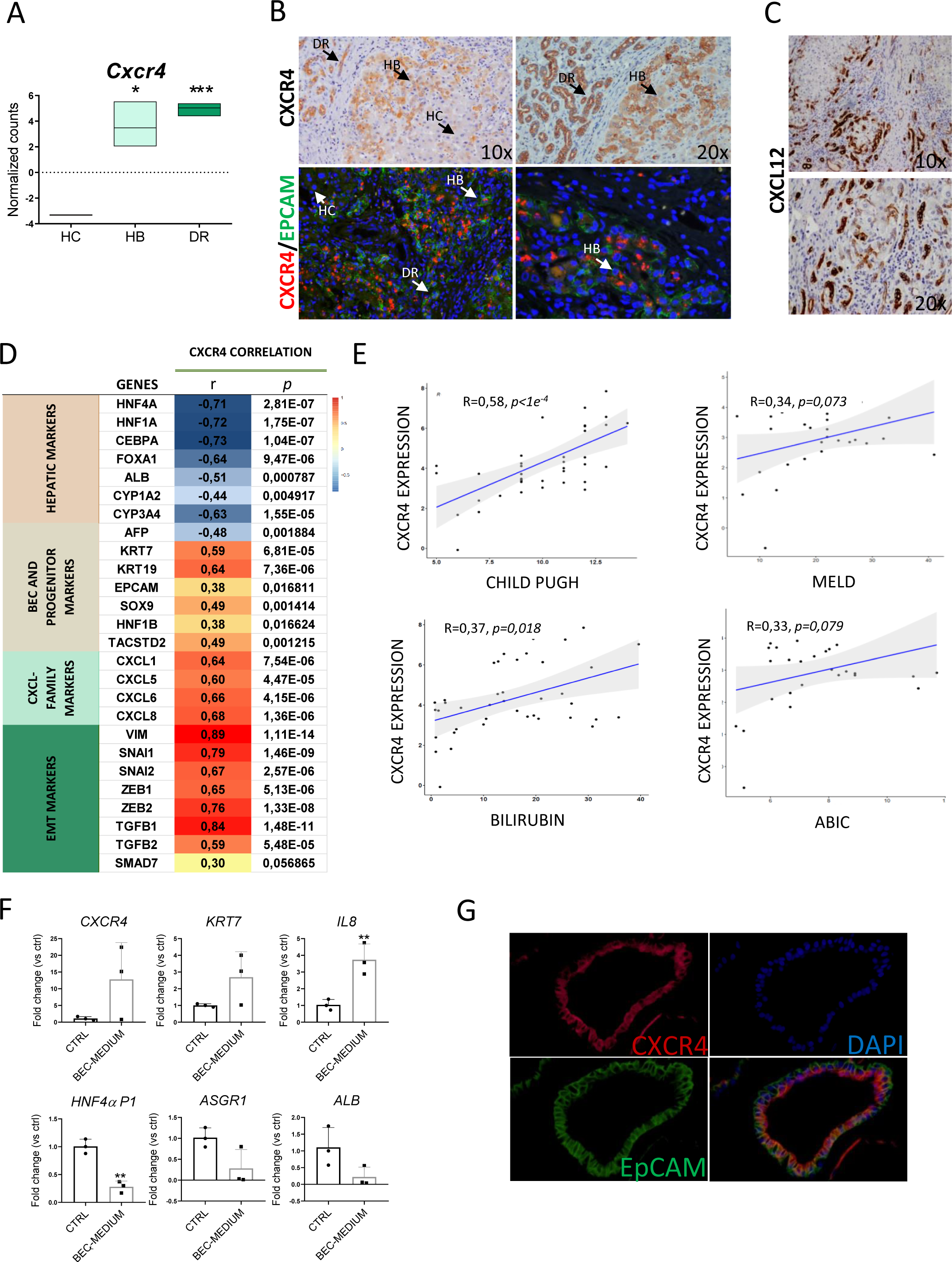
CXCR4 expression correlates with loss of hepatocyte identity and bad prognosis in AH. **(A)** Normalized counts of CXCR4 in hepatocytes, HB cells and DR cells. Significant differences are indicated as \**p*<0,05 (T-student test). **(B)** Representative staining of CXCR4 and double immunofluorescence with EPCAM and **(C)** CXCL12 on liver sections from AH patients. HC, HB and DR cells are indicated. **(D-E)** Correlation of CXCR4 expression with 4 specific gene sets: CXCL-family genes, hepatocyte markers, biliary markers, and epithelial-to-mesenchymal markers (D) and correlation with clinical parameters (E). Transcriptomic data from AH patients was used as the data set. The regression coefficient (r) and p-value of each correlation are annotated. **(F)** qPCR gene expression of AH-related human organoids incubated for 24h with the cholangiocyte organoid medium (BEC medium). Significant differences are indicated as \**p*<0,05 (T-student test). **(G)** Representative double immunofluorescence of CXCR4 and EPCAM of AH-related human organoids growth in cholangiocyte medium.

Confirming gene expression profile, immunostaining analysis showed the expression of CXCR4 and EPCAM in HB and DR cells (Fig. 4B). Moreover, CXCL12 was expressed in DR and non-parenchymal cells (Fig. 4C). Next, we evaluated the association of *CXCR4* with hepatocyte dedifferentiation in a cohort of AH patients [2]. Interestingly, *CXCR4* expression negatively correlated with hepatocyte transcription factors and albumin but showed a positive correlation with biliary and progenitor cell markers (Fig. 4D). Moreover, there was a positive correlation of *CXCR4* expression with inflammation and EMT markers (Fig. 4D). In addition, *CXCR4* gene expression correlated with ArLD disease severity scores (Fig. 4E).

We then sought to evaluate *in vitro* the role of CXCR4 on hepatocyte plasticity and dedifferentiation. First, we evaluated the effect of CXCR4 antagonist on DMSO-induced differentiated HEPARG. As TGFβ1 is known to induce hepatocyte reprogramming and up-regulate CXCR4 expression in tumor cells [25], we pretreated the cells with TGFβ1 before the incubation with the CXCR4 antagonist. As expected, TGFB-mediated up-regulation of the biliary marker *SOX9* and down-regulation of the hepatocyte markers *ALB* and the mature isoform of HNF4A (*HNF4AP1*), was partially inhibited when treated with CXCR4 antagonist AMD3100 (Fig.S1). Second, we generated AH patient-derived hepatocyte organoids. After 24h incubation with a biliary-inducing medium [20], we observed an increase of *CXCR4* expression, *KRT7 and IL8,* and a decrease of the mature subunit of HNF4A (HNF4P1), *ASGR1* and *ALB* (Fig. 4F and G). In order to evaluate the impact of CXCR4 in organoid establishment, we generated mouse organoids from CXCR4-deficient and wt hepatocytes. The efficiency of organoid formation from CXCR4-deficient hepatocytes was highly reduced, and those organoids being generated showed a markedly reduced proliferative capacity and expression of biliary and hepatocyte markers, thus indicating a role of CXCR4 in hepatocyte plasticity (Fig. 6D-F). Taken together, these results support the role of CXCR4 in hepatocyte plasticity and dedifferentiation.

### CXCR4 promotes hepatocyte dedifferentiation and chronic liver injury

To date, there are no alcohol-induced mouse models that reliably mimic the severity of AH, characterized, among other features, by the expansion of DR and hepatocyte-to-biliary reprogramming. In order to functionally study the role of *Cxcr4* expression in hepatocyte-to-biliary reprogramming and DR, we used the DDC diet mouse model, which, in contrast to ethanol-related experimental models, reproduces the main features of patients with AH. Transcriptomic profile of the DR from DDC-treated mice shows a strong similarity with the profile of the DR from AH patients as assessed by GSEA (Fig.S2A). Moreover, when compared with alcohol-based mouse models, DDC was the only model showing increased expression of Cxcr4, biliary and inflammatory markers and a reduction of hepatocyte markers, thus indicating underlining hepatocyte reprogramming (Fig. S2B) [8,20]. The presence of DR expansion, fibrosis, inflammatory recruitment and hepatocyte dedifferentiation was also confirmed at protein level in liver tissue from mice fed with DDC for 3 weeks (Fig. S2C). The up-regulation of *Cxcr4* along DDC-induced injury progression was also confirmed by qPCR, showing a similar trend as *Tgfβ1* and *Epcam* (Fig. S2D). Moreover, in order to investigate if hepatocytes expressed *Cxcr4*, primary isolated hepatocytes from healthy and DDC-treated mice were analyzed. Hepatocytes from DDC mice showed a significant increase in *Cxcr4* together with *Tgfβ, Smad3, Epcam, Snail-1, Vim*, and the fetal subunit of *Hnf4*α, *Hnf4*α*-p2,* mimicking the results observed in patients with AH (Fig. S2E).

In order to assess *in vivo* the effect of *Cxcr4* on hepatocytes, we induced *Cxcr4* overexpression by injecting intravenously AAV8-TBG-CXCR4 plasmid to C57BL6/J mice. After a 2-week washout period, mice were fed with DDC or control diet for 1 week (Fig. 5A). As shown in Fig. 5B and 5C, *Cxcr4* over-expression was confirmed in both, healthy and injured mice. Over-expression of *Cxcr4* in healthy mice induced a decreased expression of hepatocyte-specific genes (Fig. S3A) but did not induce a change in liver injury (Fig. S3B). However, mice treated with *Cxcr4* over-expression and DDC diet showed a mild non-significant increased levels of transaminases, alkaline phosphatase, and bilirubin (Fig. S4). In addition, they showed a reduced expression of mature hepatocyte genes such as *Alb*, *Cebp*α and *Cyp1a2* and increased expression of the fetal subunit of *Hnf4*α (Hnf4α p2) (Fig. 5D), suggesting that over-expression of *Cxcr4* directly promotes a loss of hepatocyte differentiation profile. Overexpression of *Cxcr4* in DDC-treated animals promoted the expansion of DR, increased the deposition of fibrosis and the periportal infiltration of neutrophils (Fig. 5E).

**Fig. 5.**
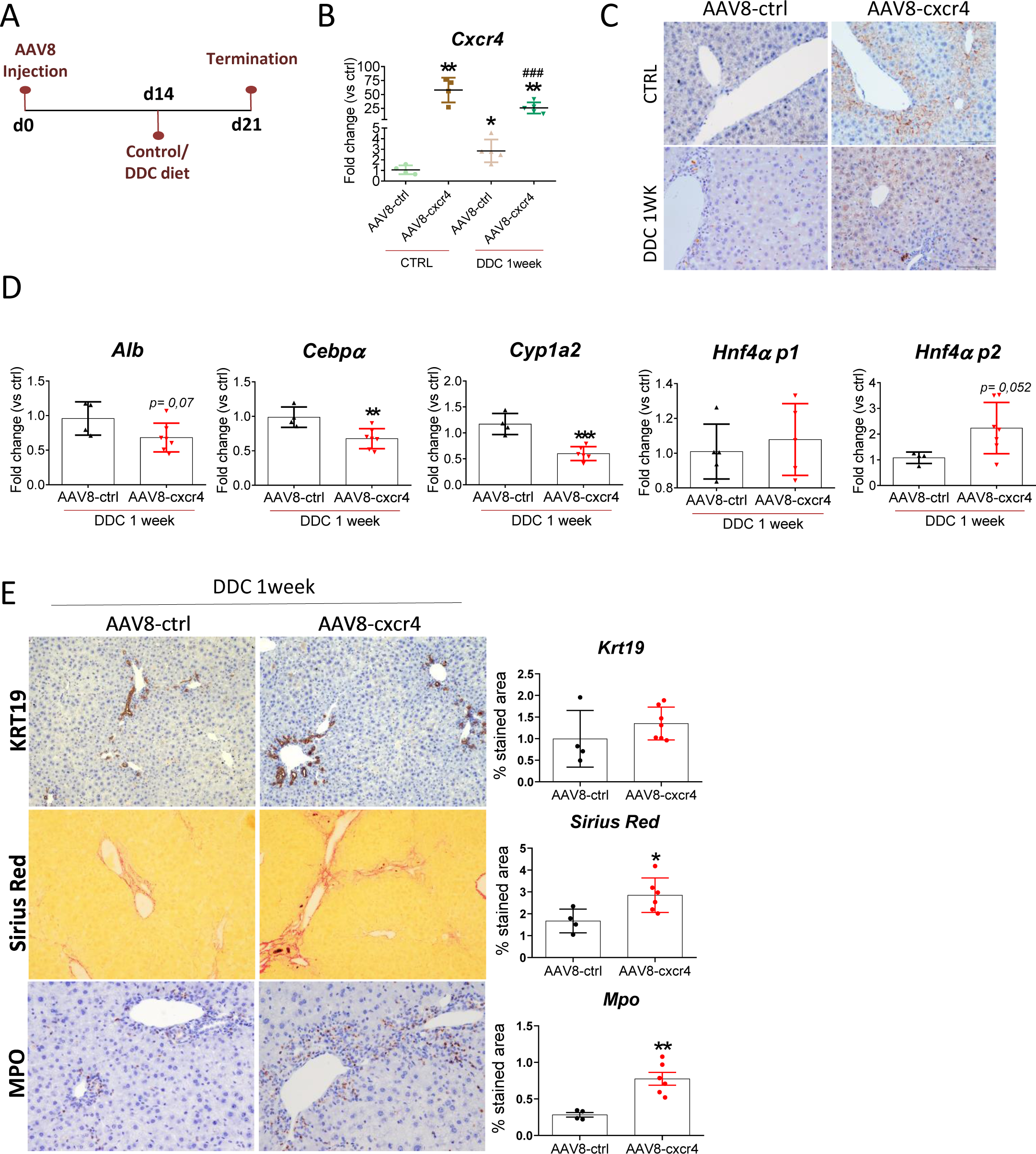
CXCR4 induces liver injury progression and loss of hepatocyte identity. **(A)** Experimental strategy to induce *Cxcr4* overexpression. **(B)** qPCR confirmation of *Cxcr4* overexpression in control (n=4) and DDC-injured mice (n=7). Gene expression is shown as Fc vs. AAV8-Ctrl. *p<0,05 compared to AAV8-Ctrl. #p<0,05 compared to AAV8-ctrl DDC 1wk (T-student test). **(C)** Representative images of CXCR4 staining in control and DDC experimental groups. **(D)** qPCR gene expression of hepatocyte-specific markers in DDC experimental groups. Data is shown as Fc vs. AAV8-Ctrl. *p<0.05 compared to Ctrl (T-student test). **(E)** Immunohistochemistry images of KRT19, MPO, and the fibrosis staining Sirius red on DDC-treated groups, AAV8-Ctrl (n=4) and AAV8-CXCR4 (n=7). Stained area quantification of both experimental groups is shown. Significant differences are indicated as \**p*<0.05 (T-student test)

**Fig. 6.**
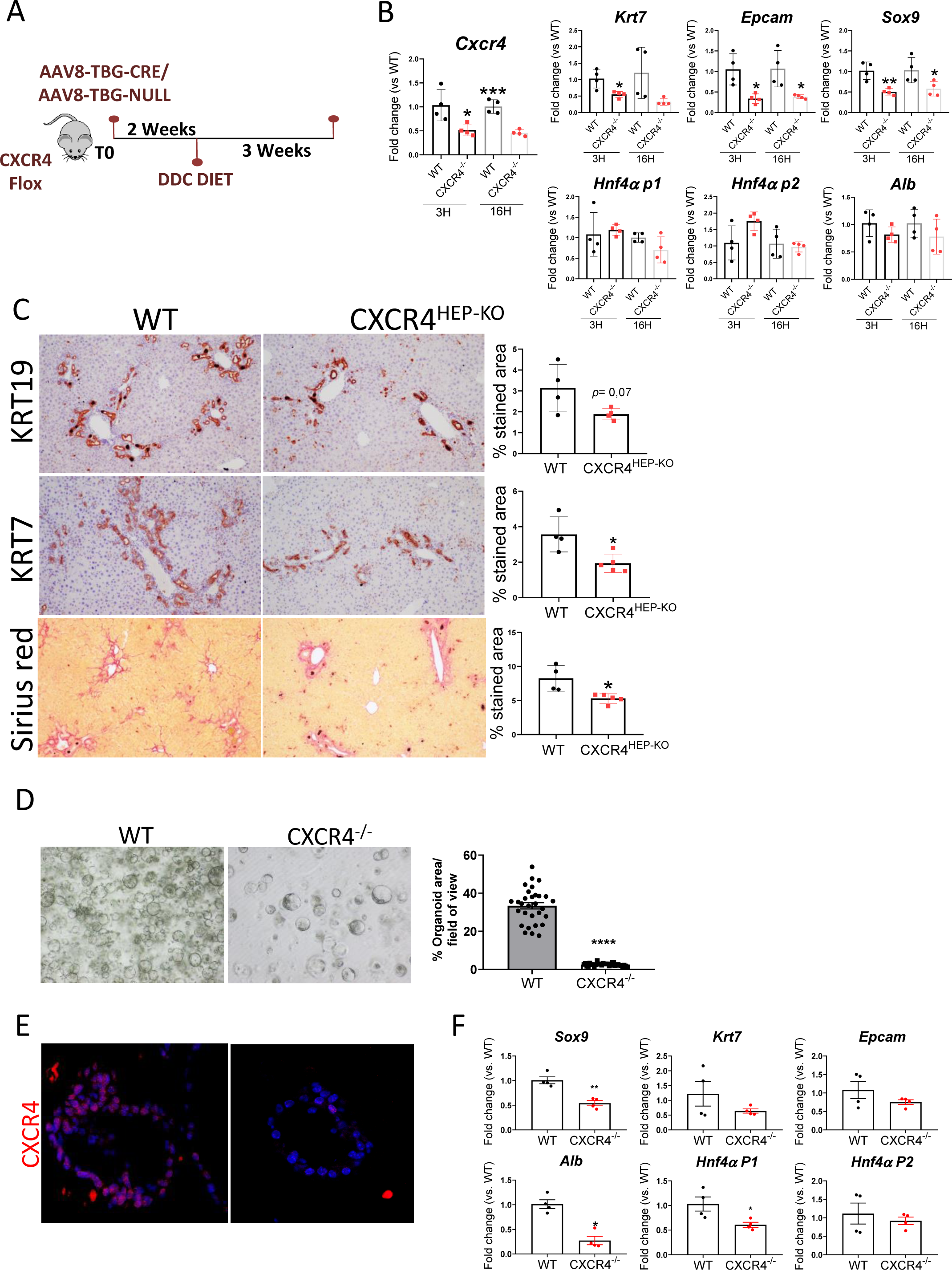
CXCR4 specific deletion in hepatocytes ameliorates hepatocyte reprogramming, DR expansion and liver fibrosis. **(A)** Experimental strategy to induce CXCR4 specific deletion in hepatocytes **(B)** qPCR of *Cxcr4, Cxcl12*, DR and hepatocyte markers on isolated primary hepatocytes from AAV8-Ctrl (WT; n=4) and AAV8-CRE treated mice (CXCR4^-/-^; n=4). **(C)** Representative images of KRT19, KRT7 and Sirius red on DDC-treated groups, AAV8-Ctrl (WT; n=4) and AAV8-CRE (CXCR4^HEP-KO^; n=5). Stained area quantification of both experimental groups is shown. Significant differences are indicated as \**p*<0.05 (T-student test) **(D)** Representative image of primary hepatocytes derived-organoids generated from AAV8-Ctrl (WT) and AAV8-CRE treated mice (KO). % Organoids area per field of view is shown. **(E)** Representative immunofluorescence of CXCR4 in primary hepatocytes derived-organoids generated from AAV8-Ctrl (WT; n=4)) and AAV8-CRE treated mice (KO; n=4). **(F)** qPCR gene expression of biliary and hepatocyte markers in mice derived-organoids. Gene expression is shown as Fc vs. WT. Significant differences are indicated as \**p*<0,05 (T-student test).

Conversely, we then investigated the effect of CXCR4 deletion by means of a CXCR4 hepatocyte specific knockout. To do so, AAV8-TBG-CRE or control plasmid were intravenously injected to CXCR4 flox mice. After a 2-week washout period, mice were fed with DDC for 3 weeks (Fig. 6A). *Cxcr4* downregulation was confirmed at gene expression level in primary isolated hepatocytes (Fig. 6B). Interestingly, knockout hepatocytes also displayed a significant decrease of the CXCR4 ligand *Cxcl12*, and the biliary markers *Krt7*, *Epcam* and *Sox9*, while no effects were found on hepatocyte markers at gene expression level (Fig. 6B). In contrast to CXCR4 overexpression, lack of CXCR4 in DDC-treated mice significantly reduced DR expansion and fibrosis deposition (Fig. 6C). Overall, these results indicate that in the context of chronic liver injury, expression of Cxcr4 in hepatocytes promotes hepatocyte loss of mature phenotype and disease progression.

### Pharmacological inhibition of the CXCL12-CXCR4 pathway reduces chronic liver injury

To assess the translational potential of targeting the CXCL12-CXCR4 pathway on chronic liver injury and hepatocyte reprogramming, we treated DDC animals with the highly specific inhibitor of Cxcr4, AMD3100. As shown in Fig. 7A, two therapeutic interventions were designed in a 3-week DDC-diet model: 1) AMD3100 administration during the last week of diet, and 2) AMD3100 administration during the last 2 weeks of diet administration (Fig. 7A).

**Fig. 7.**
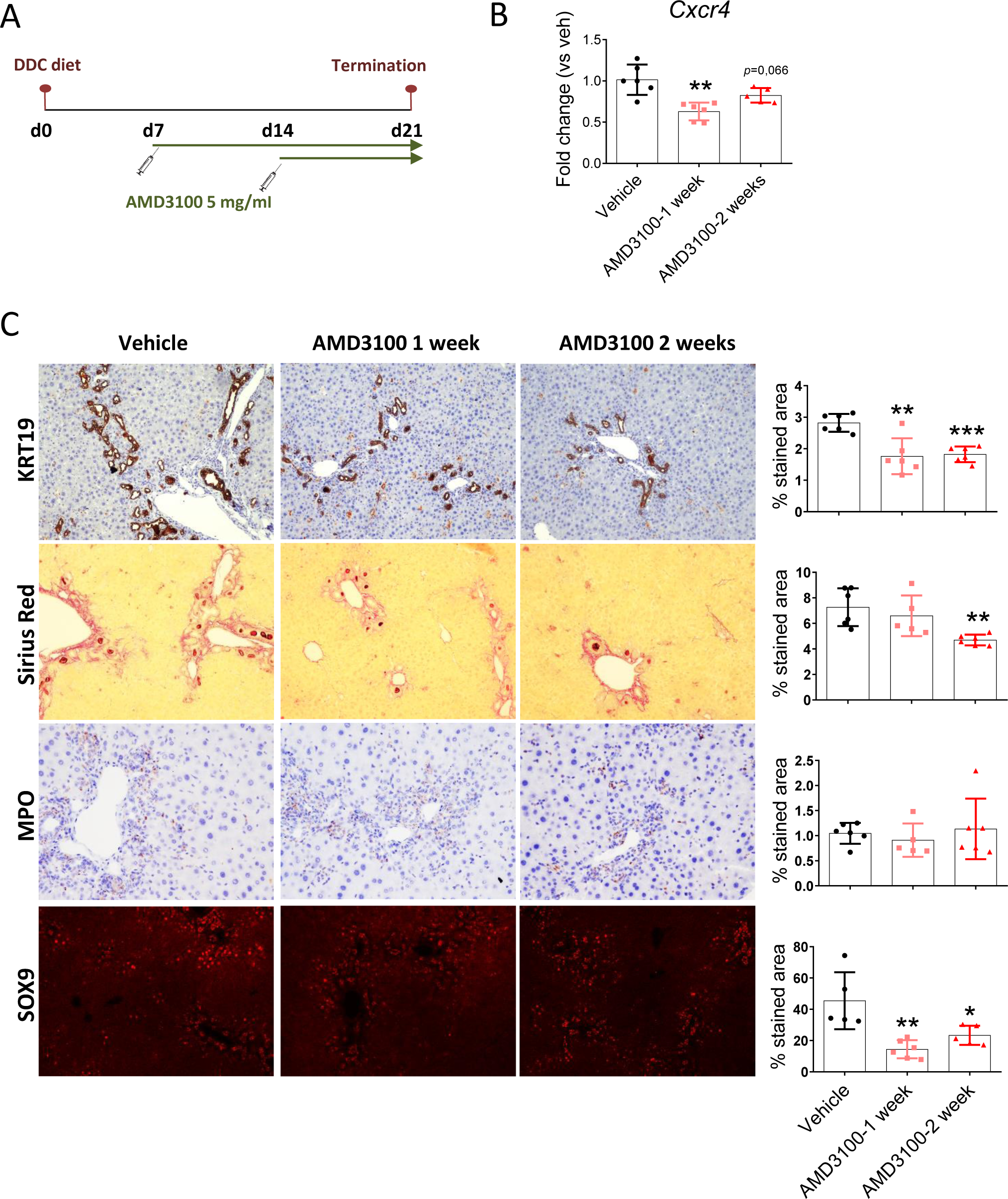
Inhibition of CXCL12/CXCR4 pathway reduces chronic liver injury. **(A)** Experimental strategy to inhibit the CXCL12/CXCR4 pathway **(B)** qPCR gene expression of *Cxcr4* in the three experimental conditions (n=6 mice per condition). Gene expression is shown as Fc vs. Vehicle **(C)** Immunohistochemistry images of KRT19, SOX9, MPO and the fibrosis staining Sirius red. Stained area quantification for each group is shown. Significant differences are indicated as \**p*<0.05 (T-student test). Images are representative of n=6 mice per condition.

Mice treated with AMD3100 showed decreased expression of *Cxcr4* (Fig. 7B), a decrease in serum bilirubin levels (Fig. S5) and no change in transaminases serum levels. In addition, treatment of mice with AMD3100 significantly reduced the extent of DR and fibrosis in both groups but had no effect on neutrophil recruitment (Fig. 7C). To evaluate the effect of Cxcl12-Cxcr4 on hepatocyte-to-biliary reprogramming, we evaluated the expression of *SOX9* in hepatocytes, which is known to be expressed in reprogrammed hepatocytes under DDC injury [26]. As shown in Fig. 7C, the number of *SOX9*-positive hepatocytes was significantly reduced in both experimental groups treated with AMD3100 as compared with the vehicle. These results demonstrate that strategies targeting Cxcl12-Cxcr4 in chronic liver diseases may prevent loss of hepatocyte identity and mitigate liver injury.

## DISCUSSION

In this study we evaluated the presence and impact of HB cells on ArLD progression. We found that HB and DR cells were increased in patients with decompensated cirrhosis and AH, however, only HB cells showed a positive correlation with disease severity and loss of hepatocyte identity. Combining LCM with bulk RNA-sequencing, we performed a comprehensive analysis of the transcriptomic profile of HB cells. We observed that HB cells present an intermediate phenotype between hepatocytes and biliary cells. Mechanistically, while overexpression of CXCR4 in hepatocytes promoted hepatocyte reprogramming, CXCR4 downregulation or its pharmacological inhibition reduced hepatocyte reprogramming and DR expansion. Overall, this study suggests that strategies limiting hepatocyte dedifferentiation and enhancing differentiation may be of interest to improve liver function in advanced ArLD.

We previously showed a positive correlation of liver progenitor cell markers with short-term mortality in patients with AH by combining histological quantification of both DR and HB cells [10]. The present study goes beyond those findings and provides independent quantification of HB cells and DR cells in patients with ArLD. Unlike DR, we showed that the presence of HB cells was a hallmark of AH. Moreover, our data suggest that hepatocyte dedifferentiation may be a specific pathogenic process taking place in ArLD concomitant to the expansion of DR. These results are not in conflict with previous reports showing a correlation of DR with disease severity [10,21,27] since we also observed that both HB cells and DR are increased in advanced stages of ArLD.

Evidence of hepatocyte plasticity have been observed in ALF and cholestatic liver diseases [2,10,13–15,22]. Previous human studies relied exclusively on histological assessment and whole liver sequencing. We specifically isolated areas of HB cells from samples of patients with AH undergoing liver transplantation [28]. This strategy enabled us to evaluate the transcriptomic profile of HB cells and to identify molecular drivers of hepatocyte-to-biliary reprogramming. In agreement with a previous study of single cell analysis of mouse reprogrammed hepatocytes [26], we found that HB cells acquired a well-characterized immature and biliary phenotype while the hepatocyte profile was, to some extent, preserved. These results may explain previous *in vivo* observations in which reprogrammed hepatocytes were able to self-renew and reacquire a mature hepatocyte phenotype [5,29,30]. Unexpectedly, we found that HB cells showed an inflammatory profile similar to that described by our group for DR cells in AH[20], and suggests that hepatocyte dedifferentiation may also be involved in inflammatory cell recruitment.

Lineage tracing studies in animal models have shown the exceptional plasticity of liver cells and have demonstrated the transdifferentiation of biliary-to-hepatocyte and hepatocyte-to-biliary processes in the context of specific animal models [29,31,32]. However, the interplay of hepatocyte reprogramming and the expansion of DR cells in human disease are still not completely understood. This study suggests that both events may take place simultaneously and may be driven by specific regulatory pathways such as CXCR4.

In the present study we found increased expression of CXCR4 in reprogrammed hepatocytes in patients with severe AH, which was associated with the loss of hepatocyte identity, stemness and disease severity. These results are in agreement with the use of CXCR4 as a biomarker to evaluate the commitment of induced pluripotent stem cells to endoderm prior to hepatocyte differentiation [33]. Additionally, the CXCR4 pathway is known to play a role in the progression of a variety of tumors [34]. Particularly, in HCC, CXCR4 expression is reportedly increased in patients with a less differentiated tumor phenotype and a cirrhotic background [35]. In this study, we provide evidence that the CXCR4 pathway directly mediates hepatocyte reprogramming in the absence of a tumor. This finding adds to the dual role of pathways such as the TGFβ or Hippo pathway, which are involved in both hepatocyte-to-biliary reprogramming and tumorigenesis [2,15],[36].

CXCR4 is involved in partial EMT in the context of liver tumor [37]. To date, there is no evidence of hepatocytes undergoing full EMT [38]. However, rather than a full EMT, a partial EMT characterized by the simultaneous expression of epithelial and mesenchymal markers may take place under injury conditions. Here we show that HB cells express a typically expressed pattern of cells undergoing partial EMT, and that this pattern correlates with CXCR4 expression, thus suggesting a potential role of CXCR4 in this process.

Targeting CXCL12/CXCR4 in experimental models attenuates liver fibrosis by reducing the activation and migration of hepatic stellate cells [39]. Moreover, in the tumor context, inhibition of CXCR4 reduces tumor malignancy [40]. Therapeutic targeting of CXCL12-CXCR4 with AMD3100 administration and genetic deletion of CXCR4 in hepatocytes in DDC-treated mice reduced hepatocyte dedifferentiation and the extent of DR and fibrosis, thus providing evidences of the potential of AMD3100 as a therapeutic strategy to improve hepatocyte differentiation and liver repair. We observed no effect on neutrophil recruitment despite the fact that this inflammatory population expresses cxcr4 and is associated with ductular reaction in alcoholic hepatitis [20]. Moreover, although the pharmacological approach may have effects other than those directly attributable to reducing hepatocyte reprograming, our results are in agreement with reports indicating that the therapeutic use of HNF4α mRNA to restore hepatocyte function reverses liver fibrosis [41]. In this regard, studies in organoids showed that hepatocyte to biliary reprogramming is associated with CXCR4 increase and mature HNF4αP1 decrease and overexpression of CXCR4 in hepatocytes promote the expression of fetal HNF4αP2 isoform. Although we find an association of CXCR4 expression with the expression of HNF4αP1/P2 isoforms, we do not have evidences of a direct mechanistic effect of CXCR4 on HNF4α regulation of hepatocyte reprogramming. Further studies will have to evaluate whether CXCR4 signaling directly regulate HNF4α expression or function.

Overall, this study uncovers the transcriptomic profile of HB cells and describes their correlation with disease progression. Moreover, it identifies the CXCR4 pathway as a new driver of hepatocyte-to-biliary reprogramming and brings to light new therapeutic strategies to promote hepatocyte maturation in ArLD.

## Abbreviations

AH: alcoholic hepatitis
ALF: acute liver failure
ArLD: alcohol-related liver disease
ASH: alcohol-related steatohepatitis
CXCR4: C-X-C motif chemokine receptor 4
DR: ductular reaction
DDC: 3,5-Diethoxycarbonyl-1,4-dihydrocollidine
DPBS: Dulbecco’s phosphate-buffered saline
EMT: epithelial-mesenchymal transition
EpCAM: Epithelial Cell Adhesion Molecule
Fc: fold change
GSEA: Gene Set Enrichment Analysis
HB: hepatobiliary cells
HCC: (hepatocellular carcinoma) Hepatocarcinoma
IPA: ingenuity pathway analysis
KRT7: cytokeratin 7
KRT19: cytokeratin 19
LCM: laser capture microdissection
NES: normalized enrichment score
qPCR: quantitative polymerase chain reaction
SOX9: SRY (Sex-Determining Region Y)-Box 9
TBG: thyroxine binding globulin
TROP2: tumor-associated calcium signal transducer 2
TGFB1: transforming growth factor beta1

## Acknowledgments

This work was performed in the Centre Esther Koplowitz. The authors wish to thank Pepa Ros for her excellent laboratory management support. We are indebted to the Genomics Unit and the Biobank Unit of the Institut d’Investigacions Biomèdiques August Pi i Sunyer (IDIBAPS) for their technical help.

---

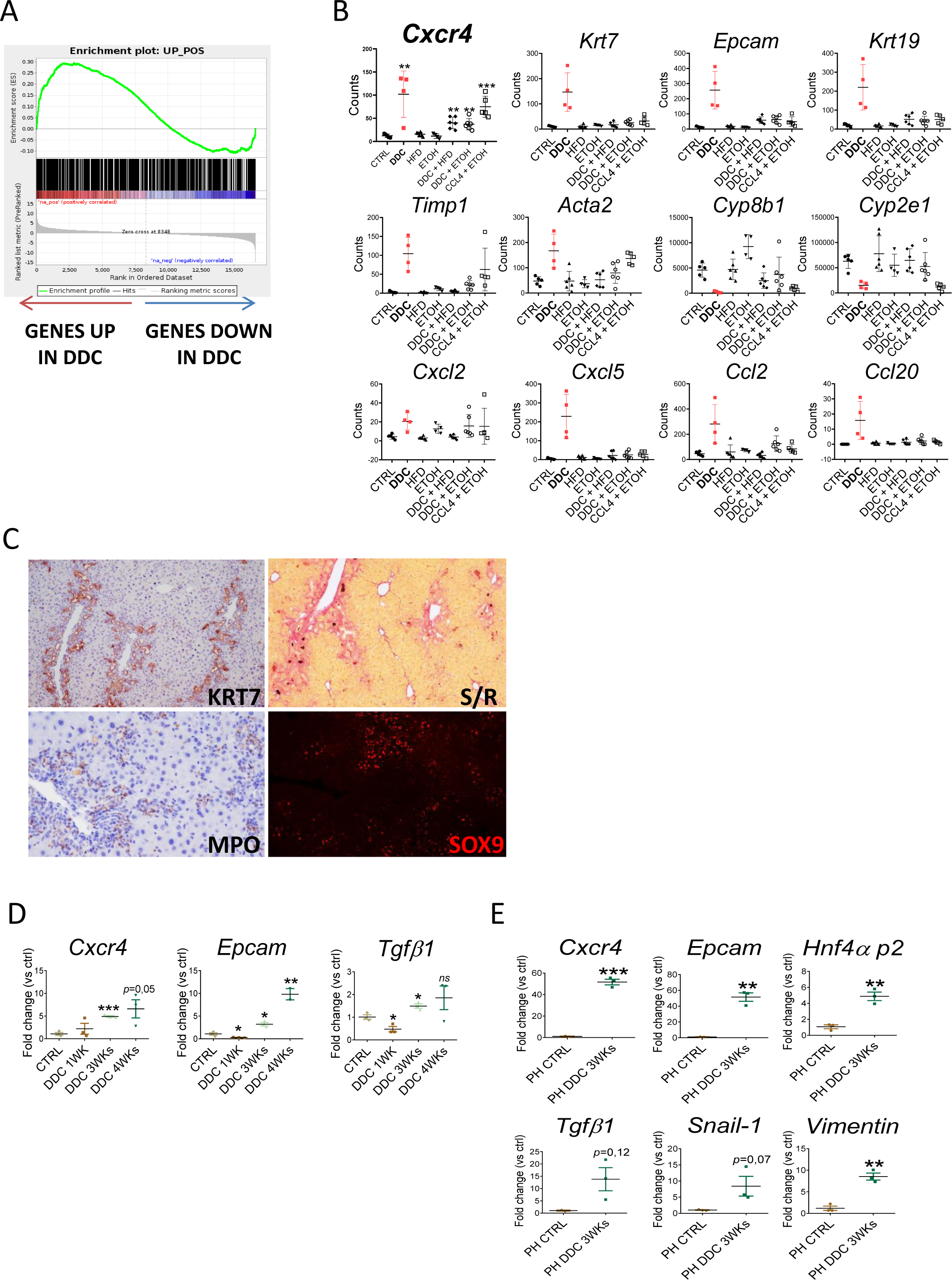

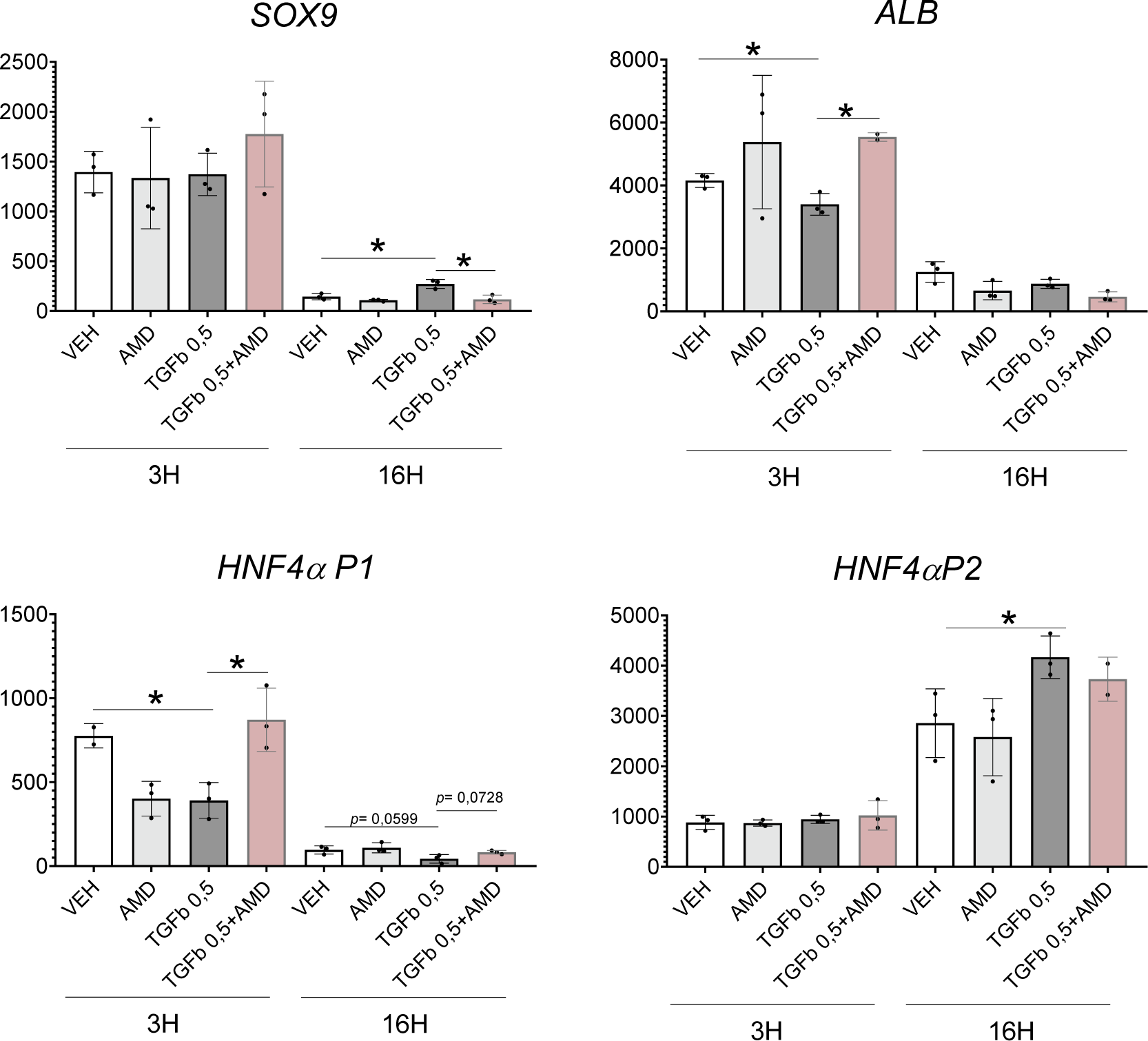

## REFERENCES

[1] Altamirano J, Miquel R, Katoonizadeh A, Abraldes JG, Duarte-Rojo A, Louvet A, et al. A histologic scoring system for prognosis of patients with alcoholic hepatitis. Gastroenterology 2014;146. https://doi.org/10.1053/j.gastro.2014.01.018.

[2] Argemi J, Latasa MU, Atkinson SR, Blokhin IO, Massey V, Gue JP, et al. Defective HNF4alpha-dependent gene expression as a driver of hepatocellular failure in alcoholic hepatitis. Nat Commun 2019. https://doi.org/10.1038/s41467-019-11004-3.

[3] Bataller R, Arab JP, Shah VH. Alcohol-Associated Hepatitis. N Engl J Med 2022;387:2436–48. https://doi.org/10.1056/NEJMRA2207599.

[4] Lucey MR, Mathurin P, Morgan TR. Alcoholic hepatitis. N Engl J Med 2009;360:2758–69. https://doi.org/10.1056/NEJMra0805786.

[5] Tanimizu N, Nishikawa Y, Ichinohe N, Akiyama H, Mitaka T. Sry HMG box protein 9-positive (Sox9+) epithelial cell adhesion molecule-negative (EpCAM-) biphenotypic cells derived from hepatocytes are involved in mouse liver regeneration. J Biol Chem 2014;289:7589–98. https://doi.org/10.1074/jbc.M113.517243.

[6] Oh SH, Swiderska-Syn M, Jewell ML, Premont RT, Diehl AM. Liver regeneration requires Yap1-TGFβ-dependent epithelial-mesenchymal transition in hepatocytes. J Hepatol 2018;69:359–67. https://doi.org/10.1016/J.JHEP.2018.05.008.

[7] Li W, Yang L, He Q, Hu C, Zhu L, Ma X, et al. A Homeostatic Arid1a-Dependent Permissive Chromatin State Licenses Hepatocyte Responsiveness to Liver-Injury-Associated YAP Signaling. Cell Stem Cell 2019;25:54–68.e5. https://doi.org/10.1016/j.stem.2019.06.008.

[8] Rodrigo-Torres D, Affò S, Coll M, Morales-Ibanez O, Millán C, Blaya D, et al. The biliary epithelium gives rise to liver progenitor cells. Hepatology 2014;60:1367–77. https://doi.org/10.1002/hep.27078.

[9] Pepe-Mooney BJ, Dill MT, Alemany A, Ordovas-Montanes J, Matsushita Y, Rao A, et al. Single-Cell Analysis of the Liver Epithelium Reveals Dynamic Heterogeneity and an Essential Role for YAP in Homeostasis and Regeneration. Cell Stem Cell 2019. https://doi.org/10.1016/j.stem.2019.04.004.

[10] Sancho-Bru P, Altamirano J, Rodrigo-Torres D, Coll M, Millán C, José Lozano J, et al. Liver progenitor cell markers correlate with liver damage and predict short-term mortality in patients with alcoholic hepatitis. Hepatology 2012;55:1931–41. https://doi.org/10.1002/hep.25614.

[11] Loft A, Alfaro AJ, Schmidt SF, Pedersen FB, Terkelsen MK, Puglia M, et al. Liver-fibrosis-activated transcriptional networks govern hepatocyte reprogramming and intra-hepatic communication. Cell Metab 2021;33:1685–1700.e9. https://doi.org/10.1016/J.CMET.2021.06.005.

[12] Roskams TA, Theise ND, Balabaud C, Bhagat G, Bhathal PS, Bioulac-Sage P, et al. Nomenclature of the finer branches of the biliary tree: Canals, ductules, and ductular reactions in human livers. Hepatology 2004. https://doi.org/10.1002/hep.20130.

[13] Hyun J, Oh SH, Premont RT, Guy CD, Berg CL, Diehl AM. Dysregulated activation of fetal liver programme in acute liver failure. Gut 2019;68:1076–87. https://doi.org/10.1136/gutjnl-2018-317603.

[14] Hyun J, Sun Z, Ahmadi AR, Bangru S, Chembazhi U V., Du K, et al. Epithelial splicing regulatory protein 2-mediated alternative splicing reprograms hepatocytes in severe alcoholic hepatitis. J Clin Invest 2020. https://doi.org/10.1172/JCI132691.

[15] Bou Saleh M, Louvet A, Ntandja-Wandji LC, Boleslawski E, Gnemmi V, Lassailly G, et al. Loss of hepatocyte identity following aberrant YAP activation: A key mechanism in alcoholic hepatitis. J Hepatol 2021;75:912–23. https://doi.org/10.1016/J.JHEP.2021.05.041.

[16] Yanger K, Zong Y, Maggs LR, Shapira SN, Maddipati R, Aiello NM, et al. Robust cellular reprogramming occurs spontaneously during liver regeneration. Genes Dev 2013;27:719–24. https://doi.org/10.1101/gad.207803.112.

[17] Tarlow BD, Pelz C, Naugler WE, Wakefield L, Wilson EM, Finegold MJ, et al. Bipotential adult liver progenitors are derived from chronically injured mature hepatocytes. Cell Stem Cell 2014;15:605–18. https://doi.org/10.1016/j.stem.2014.09.008.

[18] Yimlamai D, Christodoulou C, Galli GG, Yanger K, Pepe-Mooney B, Gurung B, et al. Hippo pathway activity influences liver cell fate. Cell 2014;157:1324–38. https://doi.org/10.1016/j.cell.2014.03.060.

[19] Fan B, Malato Y, Calvisi DF, Naqvi S, Razumilava N, Ribback S, et al. Cholangiocarcinomas can originate from hepatocytes in mice. J Clin Invest 2012;122:2911–5. https://doi.org/10.1172/JCI63212.

[20] Aguilar-Bravo B, Rodrigo-Torres D, Ariño S, Coll M, Pose E, Blaya D, et al. Ductular Reaction Cells Display an Inflammatory Profile and Recruit Neutrophils in Alcoholic Hepatitis. Hepatology 2019;69:2180–95. https://doi.org/10.1002/hep.30472.

[21] Dubuquoy L, Louvet A, Lassailly G, Truant S, Boleslawski E, Artru F, et al. Progenitor cell expansion and impaired hepatocyte regeneration in explanted livers from alcoholic hepatitis. Gut 2015;64:1949–60. https://doi.org/10.1136/gutjnl-2014-308410.

[22] Spee B, Carpino G, Schotanus BA, Katoonizadeh A, Borght S V., Gaudio E, et al. Characterisation of the liver progenitor cell niche in liver diseases: potential involvement of Wnt and Notch signalling. Gut 2010;59:247–57. https://doi.org/10.1136/gut.2009.188367.

[23] Guo F, Wang Y, Liu J, Mok SC, Xue F, Zhang W. CXCL12/CXCR4: a symbiotic bridge linking cancer cells and their stromal neighbors in oncogenic communication networks. Oncogene 2016;35:816–26. https://doi.org/10.1038/ONC.2015.139.

[24] Saiman Y, Jiao J, Fiel MI, Friedman SL, Aloman C, Bansal MB. Inhibition of the CXCL12/CXCR4 chemokine axis with AMD3100, a CXCR4 small molecule inhibitor, worsens murine hepatic injury. Hepatol Res 2015;45:794–803. https://doi.org/10.1111/HEPR.12411.

[25] Bertran E, Caja L, Navarro E, Sancho P, Mainez J, Murillo MM, et al. Role of CXCR4/SDF-1 alpha in the migratory phenotype of hepatoma cells that have undergone epithelial-mesenchymal transition in response to the transforming growth factor-beta. Cell Signal 2009;21:1595–606. https://doi.org/10.1016/J.CELLSIG.2009.06.006.

[26] Merrell AJ, Peng T, Li J, Sun K, Li B, Katsuda T, et al. Dynamic transcriptional and epigenetic changes drive cellular plasticity in the liver. Hepatology 2021:hep.31704. https://doi.org/10.1002/hep.31704.

[27] Guldiken N, Kobazi Ensari G, Lahiri P, Couchy G, Preisinger C, Liedtke C, et al. Keratin 23 is a stress-inducible marker of mouse and human ductular reaction in liver disease. J Hepatol 2016;65:552–9. https://doi.org/10.1016/j.jhep.2016.04.024.

[28] Mathurin P, Moreno C, Samuel D, Dumortier J, Salleron J, Durand F, et al. Early Liver Transplantation for Severe Alcoholic Hepatitis. N Engl J Med 2011;365:1790–800. https://doi.org/10.1056/NEJMoa1105703.

[29] Tarlow BD, Pelz C, Naugler WE, Wakefield L, Wilson EM, Finegold MJ, et al. Bipotential adult liver progenitors are derived from chronically injured mature hepatocytes. Cell Stem Cell 2014;15:605–18. https://doi.org/10.1016/j.stem.2014.09.008.

[30] Yanger K, Zong Y, Maggs LR, Shapira SN, Maddipati R, Aiello NM, et al. Robust cellular reprogramming occurs spontaneously during liver regeneration. Genes Dev 2013;27:719–24. https://doi.org/10.1101/gad.207803.112.

[31] Russell JO, Lu WY, Okabe H, Abrams M, Oertel M, Poddar M, et al. Hepatocyte-Specific β-Catenin Deletion During Severe Liver Injury Provokes Cholangiocytes to Differentiate Into Hepatocytes. Hepatology 2019;69:742–59. https://doi.org/10.1002/hep.30270.

[32] Raven A, Lu WY, Man TY, Ferreira-Gonzalez S, O’Duibhir E, Dwyer BJ, et al. Cholangiocytes act as facultative liver stem cells during impaired hepatocyte regeneration. Nature 2017;547:350–4. https://doi.org/10.1038/nature23015.

[33] Holtzinger A, Streeter PR, Sarangi F, Hillborn S, Niapour M, Ogawa S, et al. New markers for tracking endoderm induction and hepatocyte differentiation from human pluripotent stem cells. Dev 2015;142:4253–65. https://doi.org/10.1242/DEV.121020/-/DC1.

[34] Zlotnik A. New insights on the role of CXCR4 in cancer metastasis. J Pathol 2008;215:211–3. https://doi.org/10.1002/PATH.2350.

[35] Bertran E, Crosas-Molist E, Sancho P, Caja L, Lopez-Luque J, Navarro E, et al. Overactivation of the TGF-β pathway confers a mesenchymal-like phenotype and CXCR4-dependent migratory properties to liver tumor cells. Hepatology 2013;58:2032–44. https://doi.org/10.1002/HEP.26597.

[36] Liu Y, Zhuo S, Zhou Y, Ma L, Sun Z, Wu X, et al. Yap-Sox9 signaling determines hepatocyte plasticity and lineage-specific hepatocarcinogenesis. J Hepatol 2021;76:652–64. https://doi.org/10.1016/J.JHEP.2021.11.010.

[37] Malfettone A, Soukupova J, Bertran E, Crosas-Molist E, Lastra R, Fernando J, et al. Transforming growth factor-β induced plasticity causes a migratory stemness phenotype in hepatocellular carcinoma. Cancer Lett 2017;392:39–50. https://doi.org/10.1016/J.CANLET.2017.01.037.

[38] Taura K, Miura K, Iwaisako K, Ö Sterreicher CH, Kodama Y, Penz-Ö Sterreicher M, et al. Hepatocytes do not undergo epithelial-mesenchymal transition in liver fibrosis in mice. Hepatology 2010;51:1027–36. https://doi.org/10.1002/HEP.23368.

[39] Sung YC, Liu YC, Chao PH, Chang CC, Jin PR, Lin TT, et al. Combined delivery of sorafenib and a MEK inhibitor using CXCR4-targeted nanoparticles reduces hepatic fibrosis and prevents tumor development. Theranostics 2018;8:894–905. https://doi.org/10.7150/THNO.21168.

[40] Wang X, Zhang W, Ding Y, Guo X, Yuan Y, Li D. CRISPR/Cas9-mediated genome engineering of CXCR4 decreases the malignancy of hepatocellular carcinoma cells in vitro and in vivo. Oncol Rep 2017;37:3565–71. https://doi.org/10.3892/OR.2017.5601.

[41] Yang T, Poenisch M, Khanal R, Hu Q, Dai Z, Li R, et al. Therapeutic HNF4A mRNA attenuates liver fibrosis in a preclinical model. J Hepatol 2021;75:1420– 33. https://doi.org/10.1016/J.JHEP.2021.08.011.

