## Supporting information for "Hepatocyte Dedifferentiation Profiling In Alcohol-Related Liver Disease Identifies CXCR4 As A Driver Of Cell Reprogramming"

**Supplementary material and methods**

***Laser capture microdissection (LCM)***

LCM was performed in formalin-fixed paraffin embedded liver samples as previously described [1,2]. KRT7^+^ hepatocytes (light brown), KRT7^+^ DR cells (dark brown) and KRT7^-^ hepatocytes (regeneration nodules) were isolated from 3 AH patients and 1 decompensated cirrhotic patient. In order to distinguish correctly the three populations, a consecutive slide with conventional KRT7 staining was scanned and used as a guide [2]. Microdissection was performed from four patients with alcoholic hepatitis, however, due to technical problems or RNA degradation we only obtained the three cell populations from two of the patients. For those fractions missing, samples from other patients were used.

***RNA extraction, sequencing and bioinformatics analysis***

RNA extraction and sequencing was performed as previously described [1] with minor modifications. Differentially expressed genes between HB cells, DR and hepatocytes were analysed by Ingenuity Pathway Analysis (Qiagen; Red Wood, CA). Statistically significant canonical pathways and upstream regulators predicted to be inhibited (z-score< -2) or activated (z-score>2) were annotated. Pathway’s enrichment was further analyzed by Enrich R (Ma’ayan lab) database and statistically significant pathways from KEGG and GO were annotated. VolcaNoseR web app was used to visualize differentially expressed genes (up and down-regulated) between HB cells, DR and hepatocytes.

Pathway’s enrichment was evaluated using ssGSEA method included in the GSVA bioconductor package and were represented by means of balloon plot
showing scaled and centred average enrichment values by group.

***Protein interaction network***

A protein interaction network from HB gene signature was generated by string-db.org. Disconnected nodes were removed and a confidence of 0,900 was applied.

***Immunohistochemistry and Immunofluorescence***

Paraffin embedded liver sections (3 μm) from human and mouse liver samples were stained for KRT7 (1:50, DAKO, M701801), EpCAM (1:100, Dako, M080429), SOX9 (1:500, Millipore, ab5535), Hep-Par1 (ready to use, Dako, IR624), TACSTD2 (1:200, RyD, AF1112), CXCR4 (1:100, Abcam, ab124824), CXCL12 (1:100, RyD, MAB350), KRT19 (1:100, DHBS-TROMAII, AB_2133570), MPO (1:50, Abcam, ab9535), KRT7 (1:1000, Thermofisher, 15539-1-AP ), and P21(1:10, HUGO 291 CNIO).

Sections were deparaffinized and incubated in Target Retrieval Solution (Citrate Ph6, Dako), heated in a pressure cooker for 20 minutes, or rehydrated and antigen retrieved with EnVision Flex Target Retrieval Solution Low or High Ph (DAKO) in a Dako PT Link. Samples were incubated with primary antibody overnight at 4ºC. After dPBS washes, sections were incubated with secondary antibody. Diaminobenzidine (DAB, Dako) was used as a chromogen and finally, sections were counterstained with hematoxylin. For immunofluorescence staining, after incubation with secondary antibody, sections were mounted with Mounting Medium for Fluorescence with 4',6-diamidino-2-phenylindole (DAPI) (Vector Laboratories, Burlingame, CA) for nuclear staining.

***Histological quantification of hepatobiliary cells and ductular reaction cells in ArLD liver biopsies***

Liver biopsies were stained for KRT7. KRT7 staining allowed for the identification of HB cells (KRT7^+^ hepatocytes) in light brown and DR cells (KRT7^+^ DR cells) in dark brown. Consecutive images were taken in order to encompass the entire biopsies. Image quantification assessed by Image J software was performed by manual selection of the area occupied by each population. The area percenage for both populations was then calculated in relation to the total area.

***Animal models***

We used C57BL6/J mice for our study (Charles River). Male mice were used and housed under standard conditions. To induce CXCR4 overexpression, AAV8-TBG-m-CXCR4-IRES-eYFP (vector biolabs; AAV-256411) and AAV8-Ctrl plasmid (AAV8-TBG-YFP, Cat No: PV-1678) were administered intravenously in 8-week-old mice. This strategy induces the overexpression of *Cxcr4* specifically in hepatocytes, since the AAV8 virus is hepatotropic and the thyroxine binding globulin (TBG) promoter is mainly induced in hepatocytes. Briefly, AAV8 viruses were diluted in sterile Dulbecco's phosphate-buffered saline (DPBS) and administered by tail vein injection at a concentration of 2.5X10^11^ GC/ml. After a 2-week wash out period, a 0.1% 3,5-diethoxycarbonyl-1,4-dihydrocollidine (DDC) diet or control diet were given for 1 week. Conversely, to induce CXCR4 deletion, AAV8-TBG-CRE (UPV; Cat No: upv-1510C) and AAV8-Ctrl plasmid were administered intravenously in CXCR4 flox mice. After a 2-week wash out period, DDC diet was given for 3 weeks.

To inhibit the CXCL12-CXCR4 pathway, we used a 3-week DDC diet model. Specifically, the CXCR4 chemokine receptor antagonist AMD3100 (Abcam, Cambridge, United Kingdom) was resuspended in sterile DPBS and was administered intraperitoneally at 5 mg/kg dose every other day during the last week or during the last 2 weeks of the DDC diet, as shown in Fig. 6A. For both approaches, the control groups were injected with the vehicle.

All animal experiments were approved by the Ethics Committee of Animal Experimentation of the University of Barcelona and were conducted in accordance with the National Institute of Health Guide for the Care and Use of Laboratory Animals.

***Primary hepatocytes isolation***

Primary hepatocytes for *in vitro* experiments were isolated from DDC-treated mice by a three-step in situ retrograde perfusion protocol using 0.1% collagenase IV through the inferior cava vein, as previously described [3]. Pelleted hepatocytes were plated on collagen I-coated plates for gene expression analysis, or in matrigel for organoid culture:

- Pelleted hepatocytes were seeded on collagen I-coated 12-well plates (600.000/well) with William’s E medium supplemented with 10% fetal bovine serum (FBS), 2 mML-glutamine, 50 U/ml penicillin, 50mg/ml streptomycin, 1mMinsulin, 15 mM Hepes, and 50mMb-mercaptoethanol. After 4h, hepatocytes were washed 3 times with dPBS and maintained in William’s E medium supplemented with 1% FBS for 3h and 16h. RNA was then extracted for gene expression analysis.
- Primary hepatocytes were seeded on Matrigel in 24-well plates (50.000/well) as previously described [4]. Briefly, isolated hepatocytes were filtered by 70 µm filter, washed with cold AdDMEM/F12 and mixed with Matrigel. After Matrigel solidification, mouse hepatocyte medium (Hep-Medium) was added. Hep-Medium consists of AdDMEM/F12 (Thermo Fisher Scientific) supplemented with 1% HEPES, 1% GlutaMax, 1% Pen-Strep, RSPO1, B27 (minus vitamin A), 50 ng/ml EGF, 25 ng/ml HGF, 50 ng/ml FGF7 and 50 ng/ml FGF10, from Peprotech; 1µM A83-01 (Tocris); 1,25mM N-acetylcysteine, 10 nM gastrin, 3µm CHIR99021 and 10mM Nicotinamide, from Sigma; and 10 µM Rho Inhibitor γ-27632 (Calbiochem). Medium was changed every two days and 2 weeks after seeding, organoids were passaged.

***Human organoids isolation***

Organoids were obtained from liver tissue of patients with AH, as previously described [1], with minor modifications. Briefly, organoids were grown in the previously described human Hep-Medium [4] to induce a differentiated phenotype. Human Hep-Medium consist of AdDMEM/F12 (Thermo Fisher Scientific) supplemented with 1% HEPES, 1% GlutaMax, 1% Pen-Strep, RSPO1, B27 (minus vitamin A), 50 ng/ml EGF, 50 ng/ml HGF, 100 ng/ml FGF7 and 100 ng/ml FGF10, from Peprotech; 2µM A83-01 (Tocris); 1,25mM N-acetylcysteine, 10 nM gastrin, 3µm CHIR99021 and 10mM Nicotinamide, from Sigma; 20 ng/ml TGFα; and 10 µM Rho Inhibitor γ-27632 (Axon Medchem).

For Hep-organoid de-differentiation experiments, organoids were incubated for 24h with cholangiocyte-organoid medium [1]. Organoids with Hep-Medium were used as control.

***Hepatic cell line culture***

HepaRG cells were grown as previously described [5]. Briefly, HepaRG cells were cultured in William’s E medium (Sigma) supplemented with 10% FBS, 100 U/mL penicillin, 100 μg/mL streptomycin, 5 μg/mL insulin, and 50 μM hydrocortisone hemisuccinate. After 2 weeks, the medium was supplemented with 2% dimethyl sulfoxide (DMSO) and maintained for 2 more weeks. After that, HepaRG cells were detached and seeded at low confluence in the absence of DMSO and treated with 0,5 ng/ml TGFB1, 10 µg/ml AMD3100 and both together, TGFB1 and AMD3100, for 3 and 16h.

***Gene expression analysis***

Total RNA was extracted from mouse whole liver tissue using TRIzol (Life Technologies, Carlsbad, CA) and from cells using the commercial kit RNeasy Mini Kit (Qiagen) following manufacturer’s instructions. Total RNA extracted underwent quality control by Bioanalyzer 2100 (Agilent Technologies, Santa Clara, CA). Gene expression was measured using Taqman gene expression assay probes and Master Mix (Thermofisher, Waltham, MA) or SYBR Green qPCR probes and Master Mix (Applied Biosystems). mRNA levels were measured by quantitative real-time PCR (qPCR) with an ABI 7900 HT cycler (Life Technologies). Expression values were calculated based on the ∆∆Ct method. The results were expressed as 2^-∆∆Ct^.

***Statistical analysis***

Continuous variables were described as means (95% confidence interval) or medians (interquartile range). Categorical variables were described by means of counts and percentages. Comparisons between groups were performed using the Student´s *t* test or the Mann-Whitney *U* test when appropriate. Correlations between variables were evaluated using Spearman’s *rho* or Pearson’s *r*, when appropriate. The area under the receiver characteristic curve (AUROC) analysis was used to determine the best cut-off value and the accuracy (sensitivity and specificity) of continuous variables associated with mortality end points.

***Raw data information***

Microdissection raw sequencing data were deposited to the Sequence Read Archive (SRA) of NCBI under the accession number GSE199168

**Supplementary Figures**

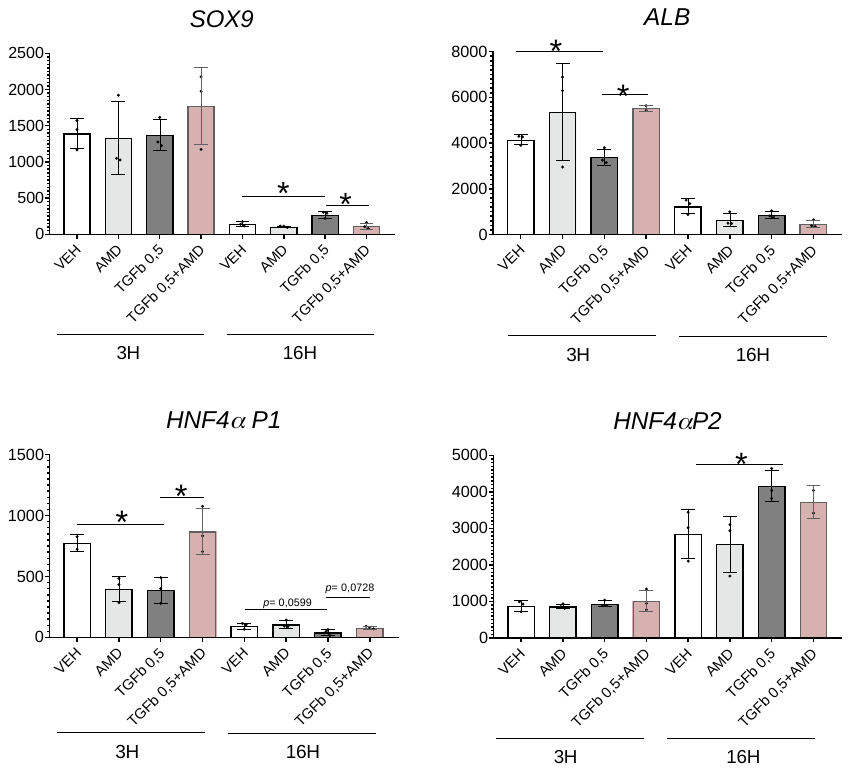
***Figure S1***

**Supplementary Fig.1. (Related to Figure 4) CXCR4 pathway inhibition reverse TGFβ-induced hepatocyte reprogramming *in vitro.* (A)** qRT-PCR analysis of differentiated HEPARG incubated with TGFB1, or TGFB1 together with CXCR4 inhibitor AMD3100 for 3h and 16h. Gene expression is shown as n=3, *p<0,05 compared to vehicle.

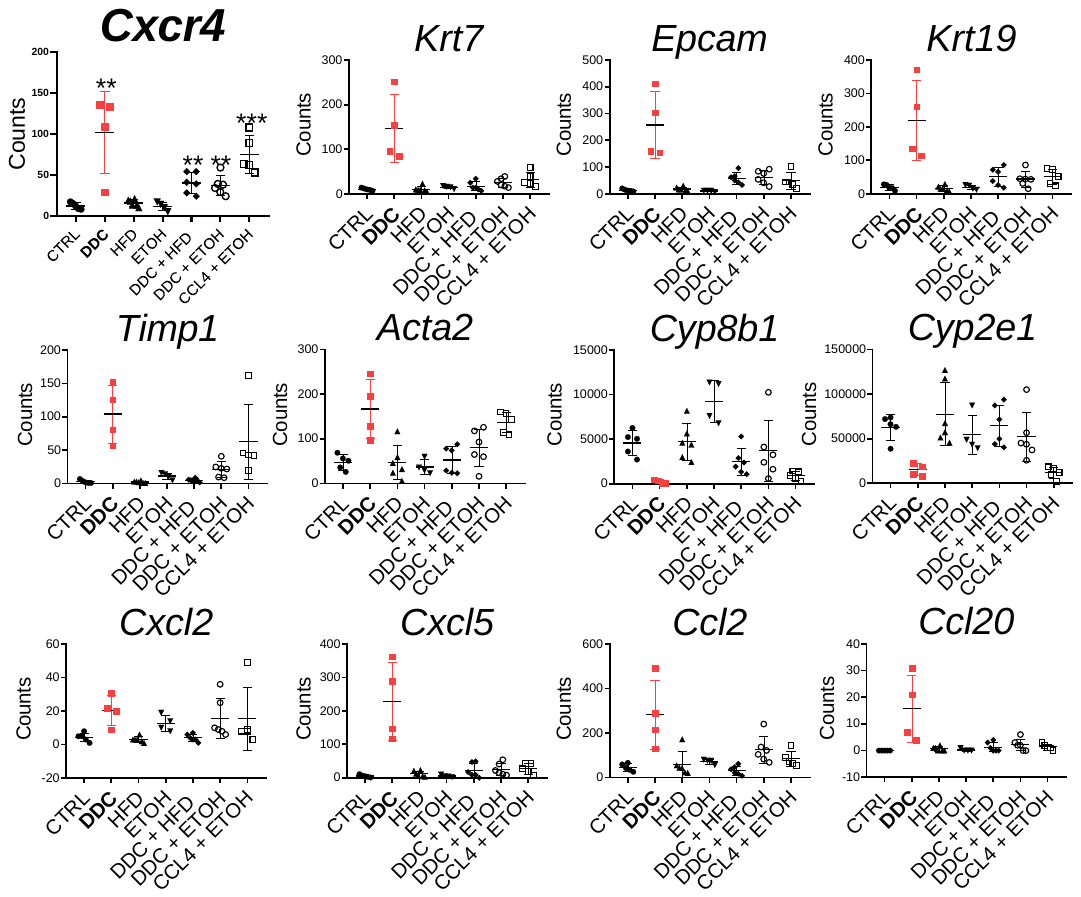
***Figure S2***

B
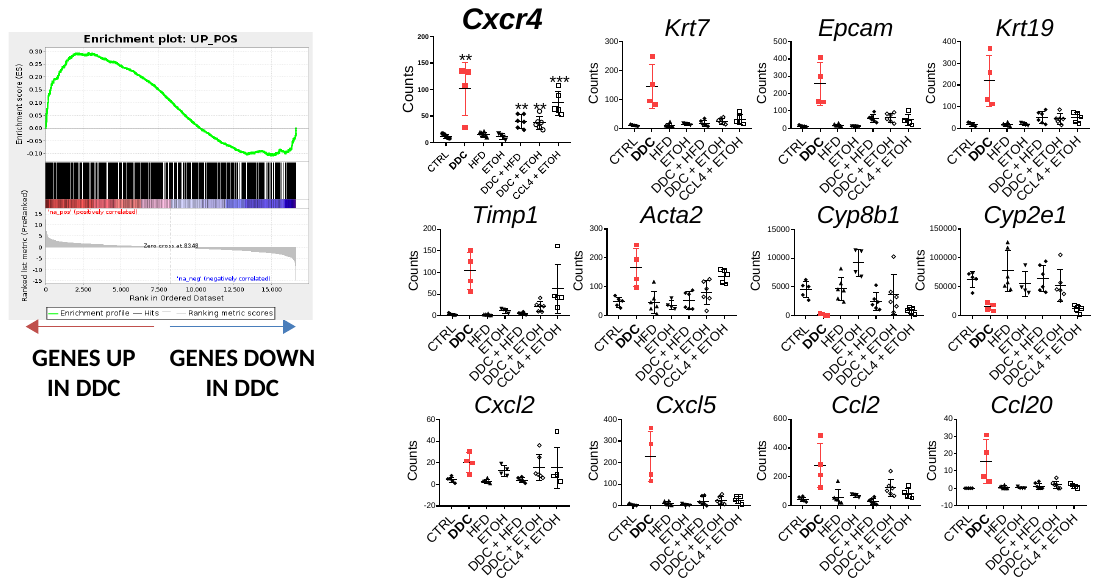

A
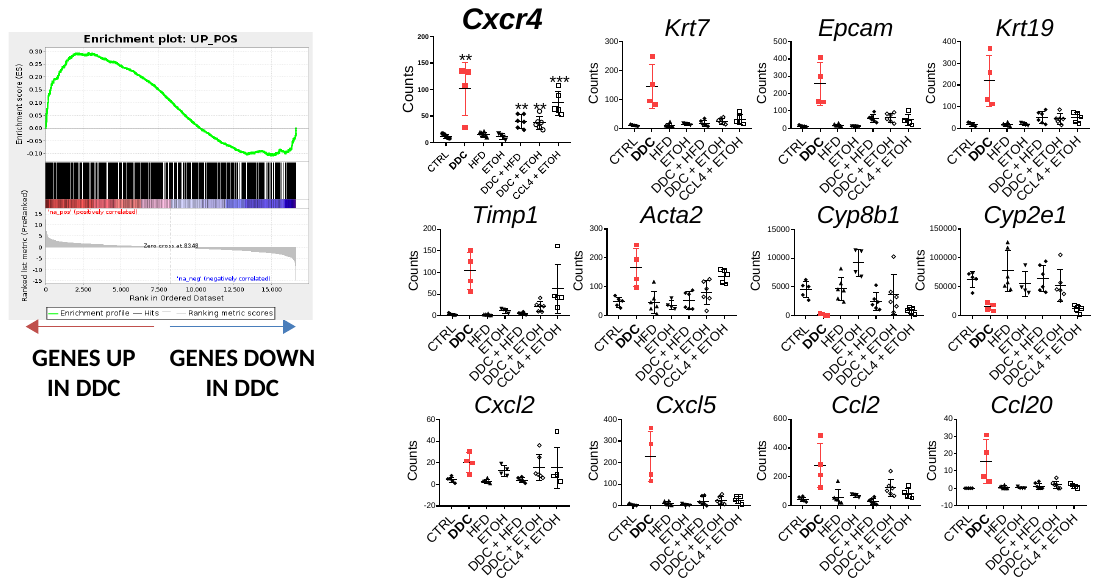

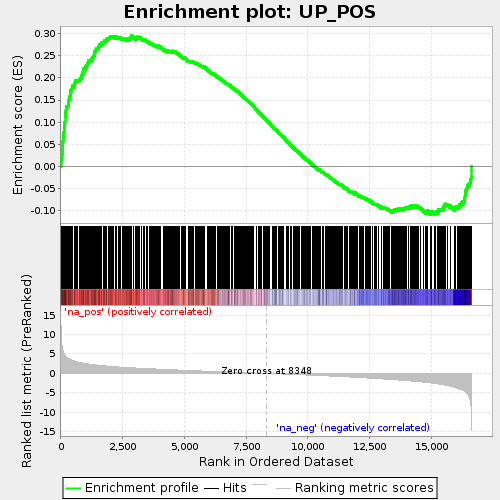

**GENES UP IN DDC**

**GENES DOWN IN DDC**

C

B

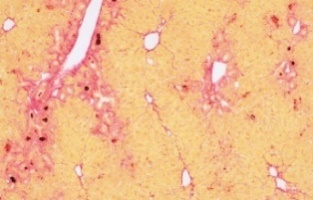

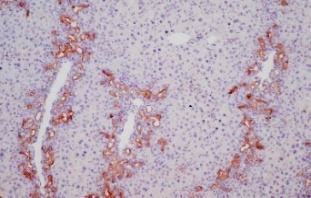

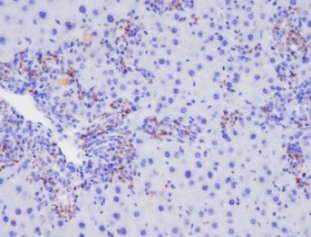

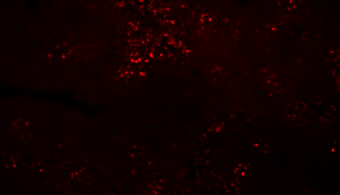

**KRT7**

**MPO**

**S/R**

**SOX9**

B

E

D

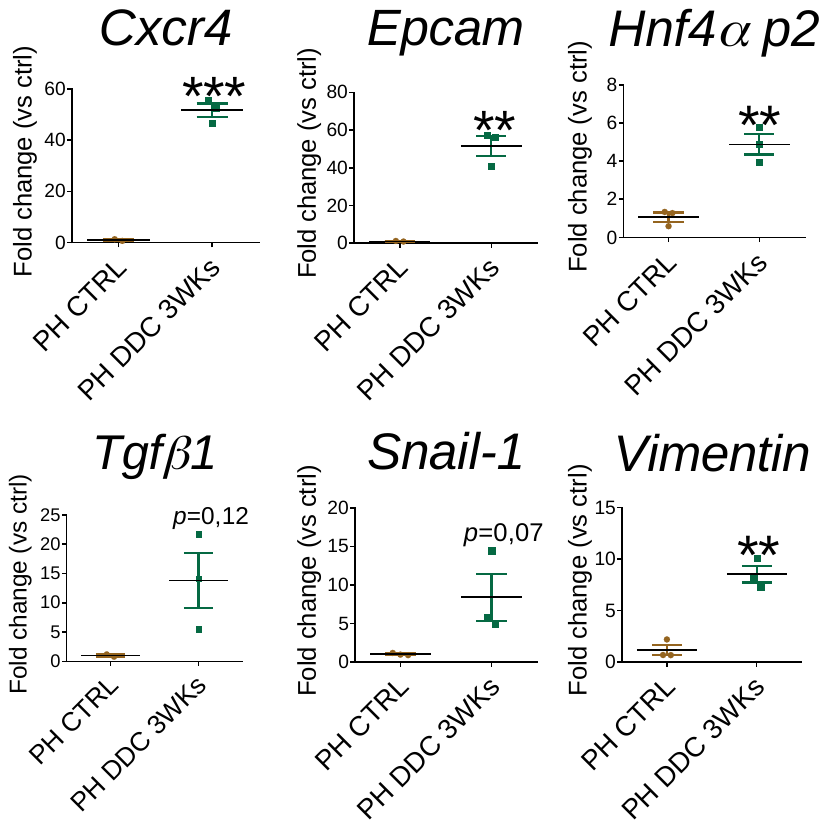

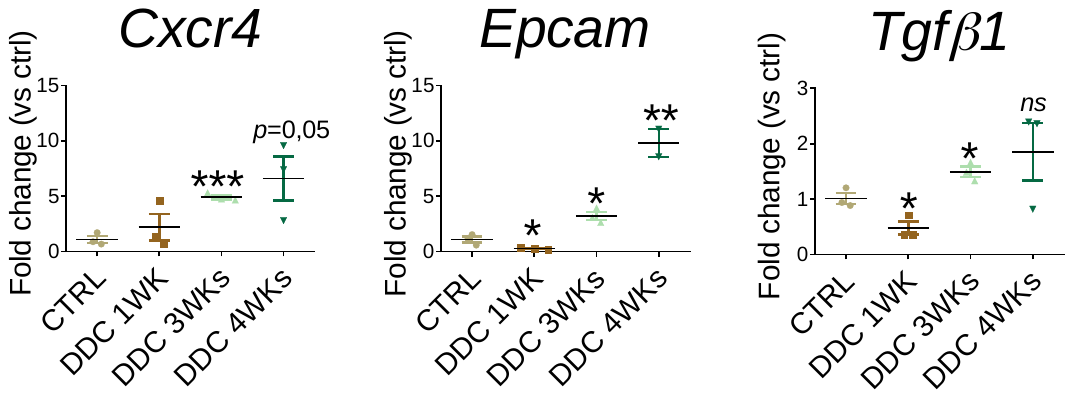

**Supplementary Fig.2 (Related to Figure 5) Similarity of the DDC mouse model with alcohol related hepatitis (AH). (A)** Gene set enrichment analysis (GSEA) between microdissected human DR transcriptome [1] and DR cells transcriptome from DDC treated mice [6]. The gene expression profile of DR cells from DDC Vs. CDE-treated mice was used as data-set. Normalized enrichment score (NES) and significance is shown. **(B)** Comparative analysis of DDC model with High Fat Diet (HFD) treated mice and alcohol-induced injury models (DDC + EtOH and CCl4 + EtOH). RNA gene expression of *Cxcr4*, ductular DR genes (*Epcam*, *Krt7* and *Krt19*), fibrosis markers (*Acta2* and *Timp1*), hepatocyte markers (*Cyp8b1* and *Cyp2e1*) and inflammatory markers (*Cxcl2*, *Cxcl5*, *Ccl2* and *Ccl20*) are shown as counts. **(C)** Histological analysis of DDC-treated mice in terms of ductular reaction (KRT7), fibrosis (Sirius red staining), neutrophil infiltration (mpo) and hepatocyte dedifferentiation (SOX9 positive hepatocytes). **(D)** Quantitative polymerase chain reaction (qPCR) gene expression of *Cxcr4*, *Tgfb1*, *Smad3* and *Epcam* in whole liver from control mice and DDC-injured mice. **(E)** Additional genes (*Snail-1*, *Vimentin* and *Hnf4αp2*) were evaluated on primary hepatocytes from control and DDC-injured mice. Gene expression is shown as fold change (Fc) vs. Ctrl. *p<0,05 compared to control mice.

***Figure S3***

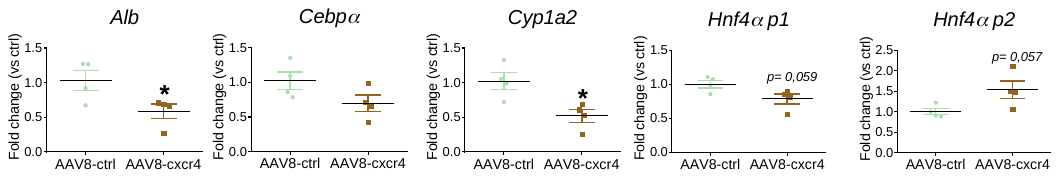

A

AAV8-ctrl

AAV8-cxcr4

**KRT19**

**Sirius Red**

B

**MPO**

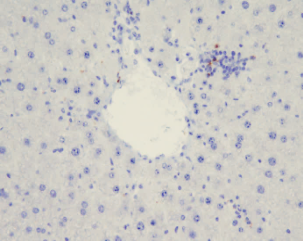

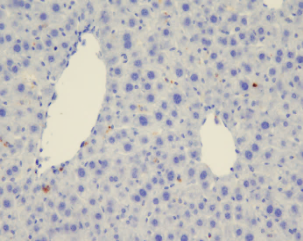

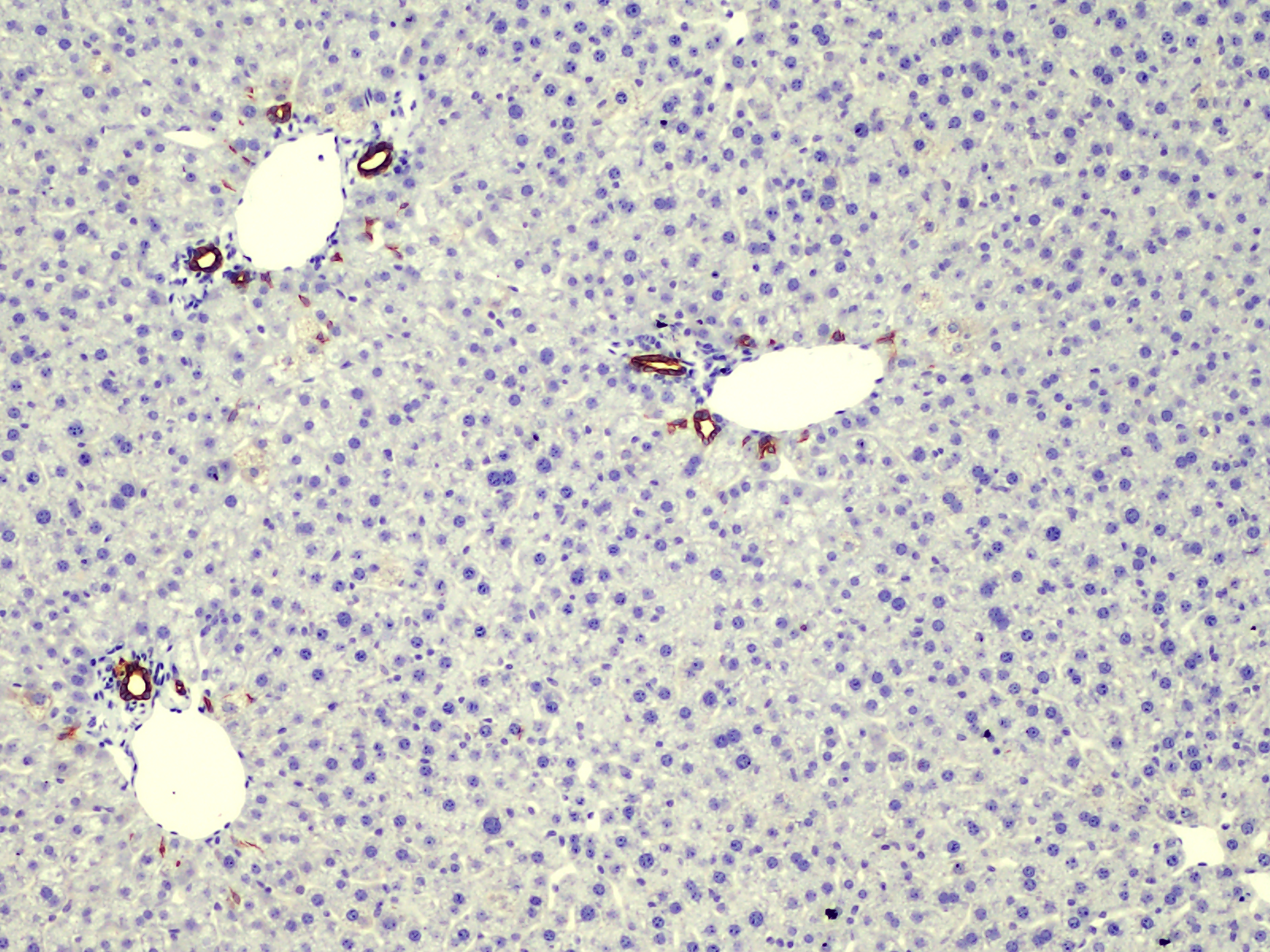

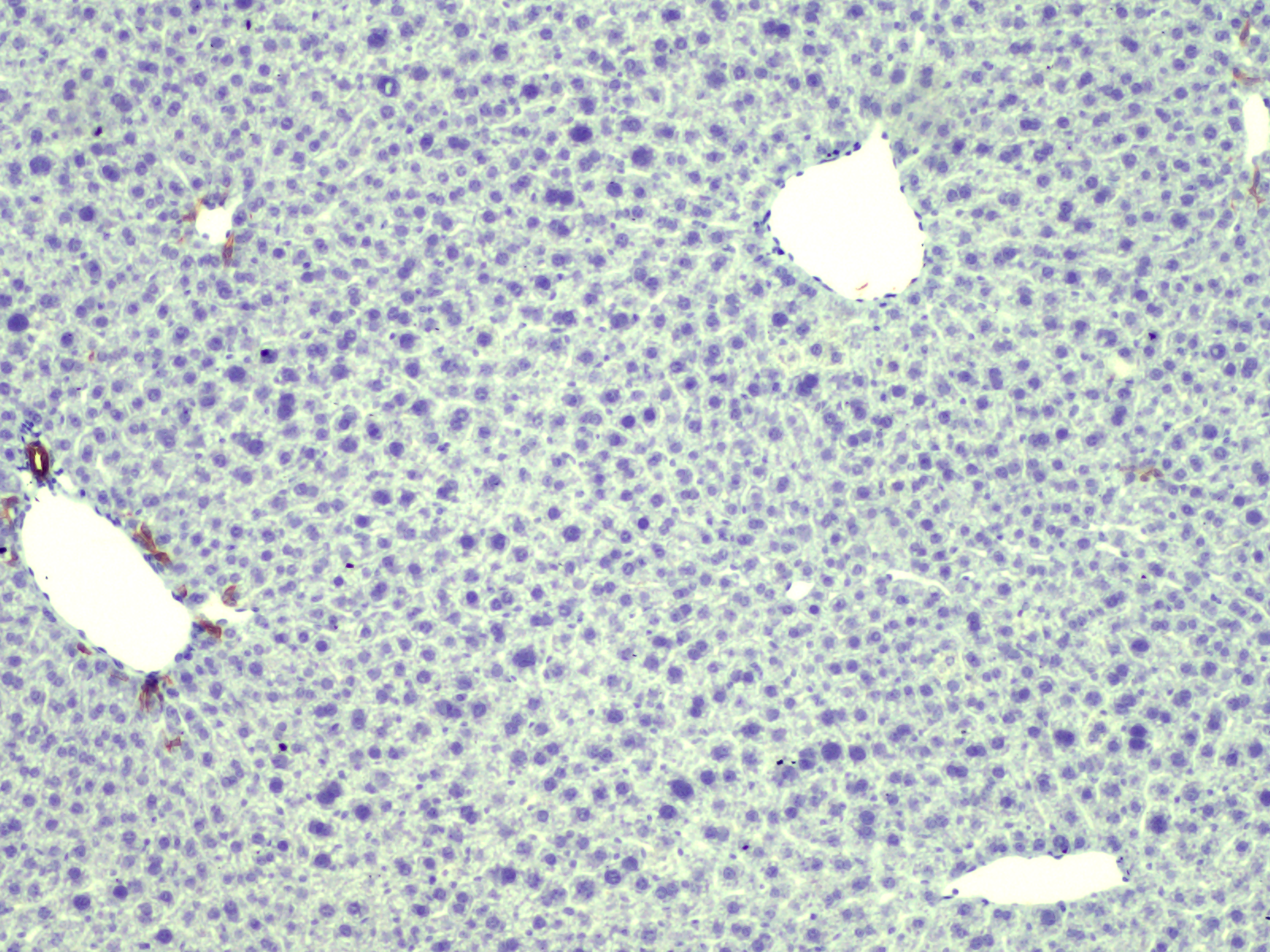

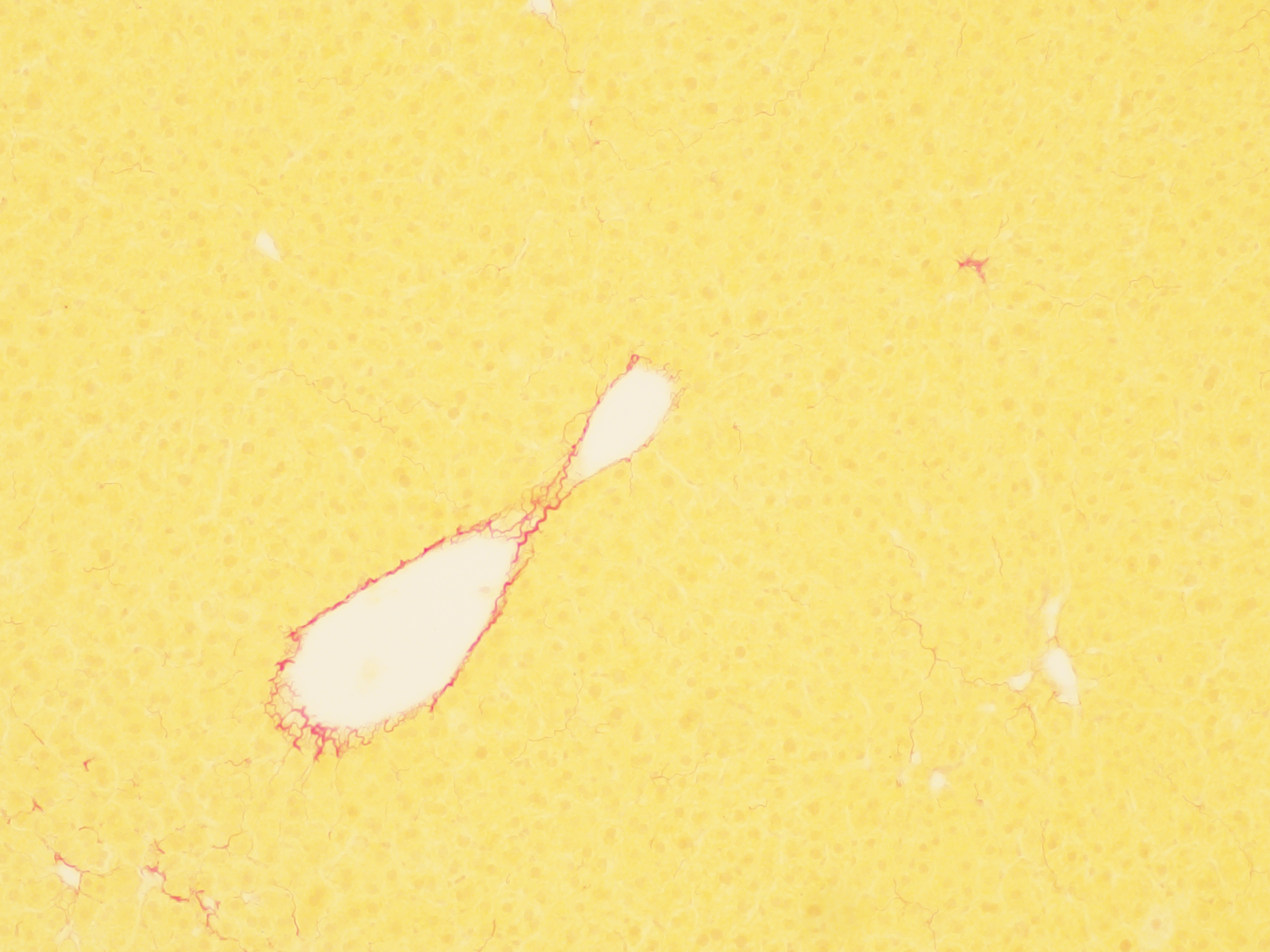

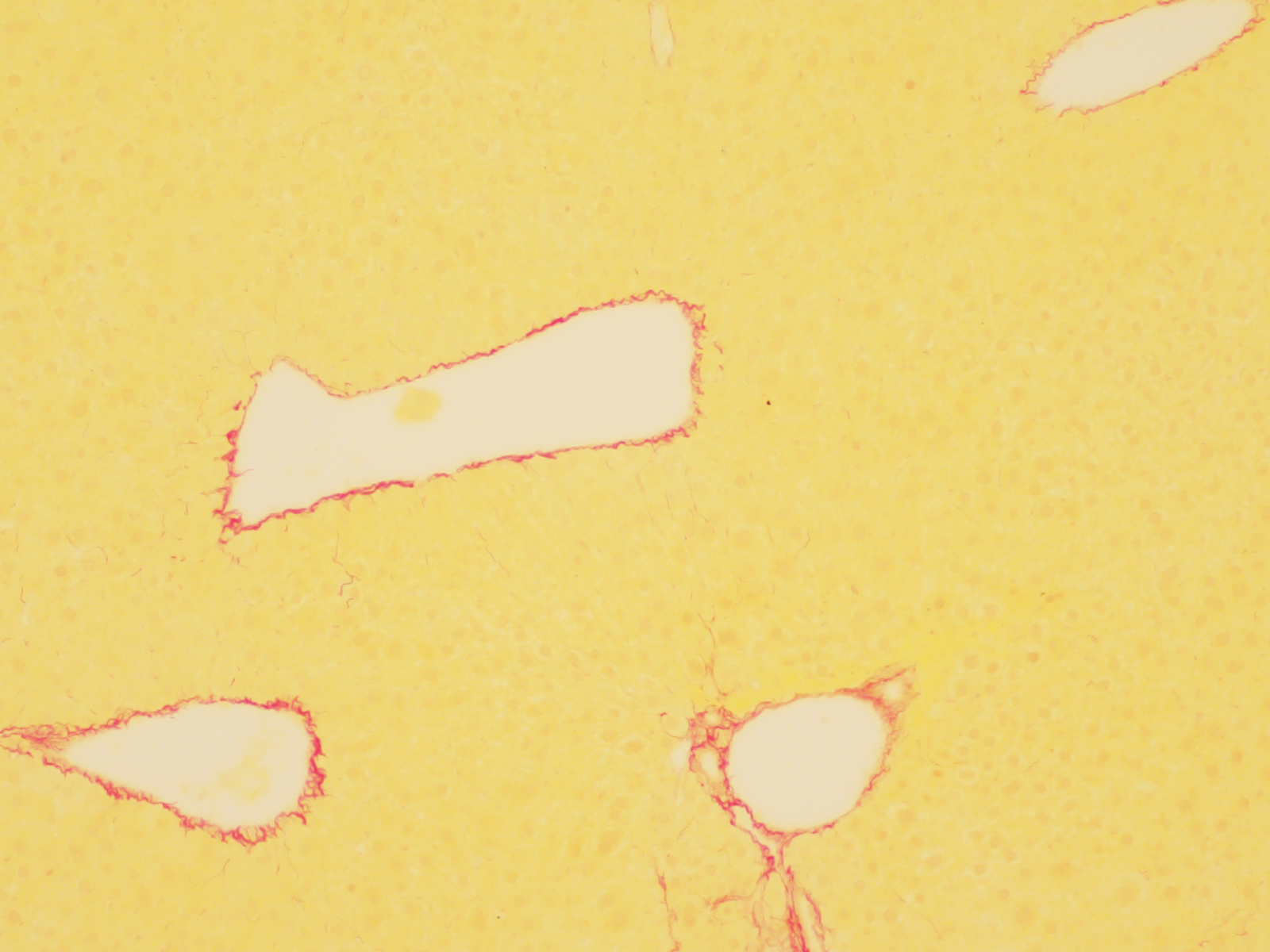

**Supplementary Fig.3. (Related to Figure5) CXCR4 overexpression in non-treated mice does not promote liver injury.** (**A)** qRT-PCR analysis of hepatocyte-specific markers in mice overexpressing CXCR4 (n=4) compared to control mice (n=4). Gene expression is shown as Fc vs control. Significant differences are indicated as **p*<0,05 (**B)** Immunohistochemistry of Krt19, Mpo, and fibrosis staining

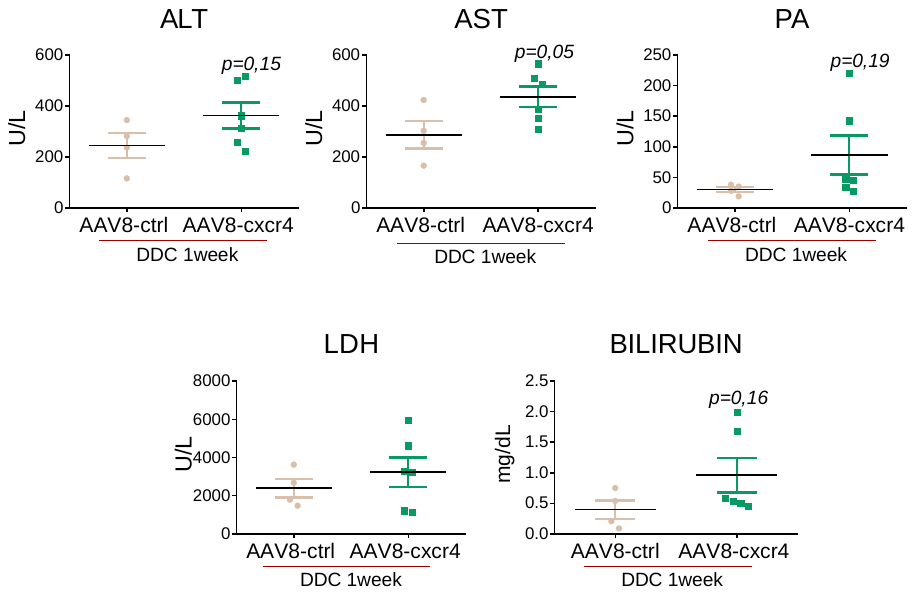
***Figure S4***

**Supplementary Fig.4. (Related to Figure5) CXCR4 overexpression in DDC-treated mice exacerbates liver injury (A)** in DDC-treated mice overexpressing CXCR4 compared to DDC-treated mice with AAV8-ctrl. *p<0,05 compared to control group.

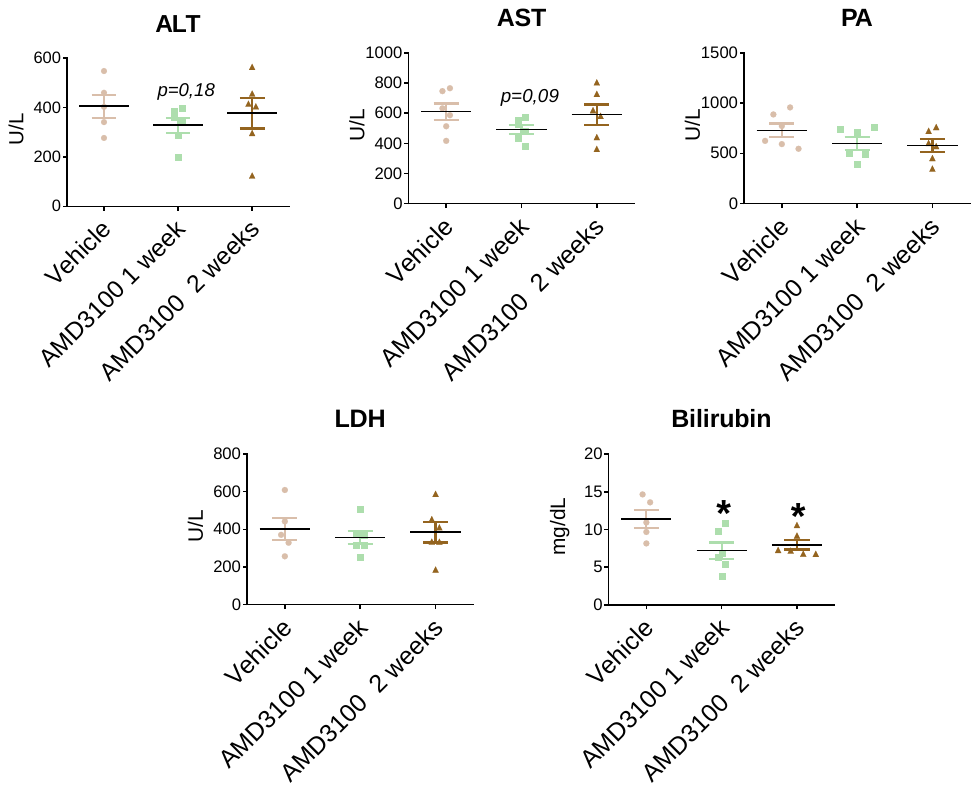
***Figure S5***

**Supplementary Fig.5. (Related to Figure6) Inhibition of CXCL12-CXCR4 pathway ameliorates liver injury. (A)** Serum levels of biochemical parameters in the three experimental conditions (vehicle, AMD3100 1week, AMD3100 2weeks). Significant differences are indicated as **p*<0,05

**Supplementary Tables**

| **Supplementary Table 1. Top 50 up-regulated and down-regulated genes in hepatobiliary cells vs. hepatocytes** | | | | | |
| --- | --- | --- | --- | --- | --- |
| **TOP 50 UP** | **Fc** | ***p*** | **TOP 50 DOWN** | **Fc** | ***p*** |
| ARNT | 767,133 | 2,54E-05 | OR6K6 | -273,64 | 9,05E-05 |
| IER5L | 374,049 | 2,74E-04 | YARS | -268,12 | 6,33E-04 |
| MMP2 | 364,15 | 6,15E-04 | GLOD5 | -221,37 | 9,85E-05 |
| STXBP1 | 330,896 | 4,33E-05 | C3orf18 | -205,63 | 2,15E-05 |
| E2F5 | 246,600 | 3,48E-04 | LIN54 | -185,39 | 4,12E-05 |
| TMEM106C | 239,260 | 9,26E-04 | SLC44A5 | -181,18 | 6,18E-04 |
| PRELP | 208,932 | 1,77E-03 | GATC | -166,07 | 7,31E-04 |
| TGFBR1 | 204,294 | 5,35E-04 | TRPV3 | -162,57 | 2,91E-04 |
| GTPBP1 | 195,846 | 9,03E-04 | SEMA3F | -156,55 | 2,33E-05 |
| DYNC2H1 | 191,865 | 8,58E-05 | DMRTA1 | -152,36 | 1,37E-03 |
| UBASH3B | 188,928 | 1,88E-03 | ZNF543 | -143,30 | 7,16E-05 |
| DCP1A | 187,909 | 3,42E-03 | LAMA3 | -141,36 | 2,60E-04 |
| CLEC4A | 180,821 | 1,25E-04 | FOXE3 | -140,69 | 2,54E-05 |
| CASP6 | 175,541 | 4,23E-03 | OR2A14 | -136,35 | 1,25E-04 |
| USP6NL | 171,538 | 3,15E-05 | CTC1 | -128,77 | 3,11E-05 |
| KCTD2 | 169,231 | 1,90E-04 | NCOA7 | -127,97 | 7,41E-05 |
| OVGP1 | 164,489 | 1,97E-03 | STOML1 | -121,97 | 1,32E-04 |
| MTX3 | 163,480 | 1,93E-03 | TPRN | -121,41 | 2,90E-03 |
| SLC25A32 | 153,417 | 2,82E-04 | PSPH | -120,44 | 6,19E-04 |
| UPF3B | 146,556 | 9,51E-05 | FBLN7 | -119,14 | 3,71E-05 |
| CCDC146 | 142,962 | 1,23E-04 | SORCS2 | -112,79 | 1,86E-04 |
| C3AR1 | 139,747 | 6,28E-04 | GBGT1 | -112,62 | 4,61E-04 |
| ANKRD13A | 139,260 | 8,75E-04 | P4HTM | -111,59 | 3,65E-04 |
| OR1N1 | 138,949 | 3,05E-04 | HOMER3 | -107,59 | 4,96E-04 |
| USP10 | 137,232 | 4,16E-05 | WDR74 | -104,75 | 8,55E-03 |
| SLC1A5 | 133,541 | 1,66E-04 | DEF8 | -93,27 | 2,63E-03 |
| MLH3 | 130,384 | 1,36E-04 | TESK1 | -91,72 | 9,51E-03 |
| CKAP5 | 126,483 | 1,60E-03 | BCL2L12 | -87,13 | 2,93E-03 |
| GUCY1A3 | 124,476 | 6,10E-03 | XPNPEP2 | -84,45 | 1,63E-01 |
| MTF2 | 117,998 | 7,40E-03 | HDAC11 | -81,68 | 3,87E-05 |
| NAT1 | 116,663 | 8,91E-04 | S100A13 | -77,75 | 1,01E-02 |
| POLDIP3 | 112,402 | 1,12E-03 | DEPTOR | -77,33 | 9,21E-02 |
| CXCR4 | 112,309 | 5,59E-03 | DPM3 | -76,29 | 3,41E-04 |
| DTWD2 | 109,987 | 2,24E-02 | GPR153 | -74,67 | 1,55E-02 |
| ZC3H8 | 109,519 | 1,44E-03 | MED31 | -74,07 | 6,07E-04 |
| PDE4DIP | 107,605 | 5,95E-03 | LMF1 | -73,48 | 7,53E-02 |
| SIN3A | 106,647 | 2,66E-04 | LTN1 | -72,58 | 9,60E-04 |
| CCDC7 | 99,927 | 3,33E-03 | PDCD7 | -69,63 | 1,09E-02 |
| TSPAN7 | 99,725 | 4,55E-03 | CCDC107 | -69,16 | 2,85E-02 |
| CPEB2 | 96,579 | 7,27E-03 | CMTR1 | -69,04 | 1,03E-01 |
| PEX6 | 94,802 | 1,82E-03 | JOSD1 | -65,18 | 1,56E-02 |
| NIN | 94,527 | 3,06E-03 | TYMS | -64,85 | 1,55E-02 |
| UBE3D | 92,708 | 4,34E-04 | ASMTL | -64,21 | 9,03E-02 |
| ZNRF3 | 91,984 | 2,98E-03 | MN1 | -62,25 | 8,09E-05 |
| EXOC2 | 90,851 | 1,20E-02 | OR6C1 | -61,75 | 8,43E-04 |
| PFKFB3 | 90,722 | 9,86E-03 | SETMAR | -61,19 | 5,24E-02 |
| OR8K3 | 87,927 | 1,08E-04 | ZNF324 | -60,51 | 1,15E-02 |
| SHPRH | 86,937 | 2,25 E-05 | POLR3A | -60,37 | 8,40E-02 |
| CXCL6 | 86,866 | 0,092274 | LRRC46 | -59,33 | 8,29E-03 |
| GLTSCR1 | 86,492 | 6,26E-05 | ZNF212 | -58,93 | 1,75E-03 |

| **Supplementary Table 2. Top 50 up-regulated and down-regulated genes in hepatobiliary cells vs. ductular reaction cells** | | | | | |
| --- | --- | --- | --- | --- | --- |
| **TOP 50 UP** | **Fc** | ***p*** | **TOP 50 DOWN** | **Fc** | ***p*** |
| MASP2 | 416,68 | 1,35E-05 | GULP1 | -611,64 | 1,60E-05 |
| SLC27A2 | 279,04 | 8,47E-06 | LAMA3 | -599,76 | 3,74E-06 |
| CHRNA4 | 272,33 | 1,45E-05 | MUC6 | -543,38 | 1,85E-03 |
| PRPS1 | 256,30 | 1,21E-05 | ZNF254 | -449,99 | 4,88E-05 |
| GBP7 | 216,81 | 2,12E-05 | GRHL2 | -395,50 | 5,15E-05 |
| CTH | 202,72 | 4,60E-05 | CTNND2 | -359,26 | 4,47E-05 |
| HIST1H3J | 176,51 | 9,36E-03 | C9orf3 | -330,30 | 4,18E-05 |
| ENPP7 | 172,22 | 1,46E-05 | B4GALNT3 | -325,77 | 6,07E-06 |
| PGLYRP2 | 163,84 | 7,69E-04 | SLC5A1 | -305,49 | 2,89E-05 |
| RORC | 148,14 | 2,56E-04 | RFX3 | -285,23 | 1,48E-05 |
| F10 | 144,28 | 3,82E-03 | EVC2 | -241,71 | 1,92E-05 |
| APOF | 143,34 | 1,25E-02 | MTCL1 | -199,80 | 2,63E-03 |
| BCAS2 | 128,91 | 1,01E-02 | ZNF543 | -191,80 | 3,83E-04 |
| PFKFB1 | 127,98 | 6,69E-04 | VEPH1 | -190,08 | 1,45E-04 |
| RAPGEF4 | 118,46 | 4,74E-04 | ZNF836 | -189,93 | 1,62E-03 |
| SLC47A1 | 110,32 | 3,32E-04 | ITGB8 | -182,44 | 1,99E-04 |
| TENM1 | 107,64 | 6,14E-04 | HOMER3 | -181,98 | 1,69E-04 |
| F7 | 107,63 | 1,77E-03 | WNK2 | -164,53 | 2,91E-04 |
| MOGAT2 | 106,80 | 5,83E-03 | PTGFR | -163,85 | 1,25E-04 |
| SNX4 | 105,89 | 8,44E-03 | DENND6B | -152,07 | 2,48E-04 |
| SERPIND1 | 104,19 | 2,01E-02 | RAB25 | -144,39 | 9,13E-05 |
| TUBE1 | 104,04 | 8,75E-03 | PRSS16 | -140,33 | 3,85E-05 |
| MRPS28 | 98,48 | 2,18E-03 | TAF1C | -136,95 | 1,68E-03 |
| ETNK2 | 94,41 | 3,43E-03 | SCTR | -135,60 | 2,87E-04 |
| AKR1C4 | 94,11 | 8,89E-03 | C3orf52 | -135,22 | 1,65E-05 |
| HPR | 93,50 | 2,13E-02 | MMP7 | -131,42 | 1,27E-02 |
| RTP3 | 90,30 | 7,27E-05 | LDOC1 | -124,77 | 1,45E-02 |
| KLKB1 | 86,71 | 5,34E-03 | ITPR3 | -119,75 | 2,63E-05 |
| ENO3 | 85,51 | 7,45E-03 | PPP1R13L | -119,01 | 2,85E-03 |
| TMA16 | 83,81 | 1,85E-04 | KCNJ16 | -114,67 | 5,49E-03 |
| KCTD18 | 83,18 | 9,49E-03 | PDE5A | -114,20 | 2,28E-02 |
| PBLD | 81,62 | 9,52E-03 | TRPV6 | -114,10 | 1,35E-02 |
| HFE2 | 81,30 | 4,34E-05 | IGFBP6 | -109,95 | 5,74E-05 |
| GYS2 | 79,88 | 1,57E-02 | TCTN2 | -108,50 | 7,78E-05 |
| CETN2 | 79,09 | 6,18E-04 | TRIM2 | -106,04 | 5,32E-04 |
| MTTP | 78,55 | 1,26E-04 | FAM3B | -104,87 | 5,48E-03 |
| MRPL10 | 77,98 | 2,64E-04 | PCNXL2 | -101,38 | 6,88E-03 |
| PEX19 | 77,17 | 2,05E-04 | CFAP70 | -98,94 | 1,41E-03 |
| TMEM82 | 76,81 | 1,04E-02 | SIRT6 | -97,20 | 3,95E-05 |
| OVGP1 | 76,53 | 1,57E-03 | CLIC6 | -95,92 | 1,20E-03 |
| VPS33A | 76,07 | 1,34E-04 | PROM1 | -95,73 | 2,46E-02 |
| RDH16 | 73,81 | 4,41E-03 | ARHGEF10 | -93,17 | 1,14E-03 |
| IL1RN | 73,01 | 7,39E-04 | SLC29A2 | -91,75 | 1,62E-03 |
| PC | 72,34 | 3,50E-04 | GALNT7 | -91,71 | 2,23E-04 |
| SMOC1 | 69,81 | 1,72E-02 | ZHX2 | -86,94 | 1,68E-02 |
| MOSPD1 | 68,75 | 5,57E-03 | GABRE | -86,77 | 2,34E-02 |
| PSAT1 | 68,11 | 1,53E-04 | B3GALT4 | -86,22 | 6,99E-05 |
| GNAO1 | 65,04 | 6,12E-03 | AQP1 | -85,05 | 5,79E-03 |
| SLC28A1 | 63,14 | 3,15E-02 | LAMC2 | -84,37 | 4,81E-03 |
| PRSS3 | 62,93 | 3,62E-04 | EHF | -83,99 | 2,12E-03 |

- Supplementary [Tables S3 and S4.xlsx](Tables%20S3%20and%20S4.xlsx)

| **Supplementary Table 5. Differential gene expression of hepatic and biliary-specific genes in the three microdissected populations** | | | | |
| --- | --- | --- | --- | --- |
| **GENE** | **Fc HB vs HC** | ***p* HB vs HC** | **Fc HB vs DR** | ***P* HB vs DR** |
| AHSG | -2,00 | 0,305 | 10,78 | 0,008 |
| ALB | -1,53 | 0,604 | 8,13 | 0,012 |
| ASGR1 | -1,46 | 0,440 | 2,84 | 0,098 |
| HNF4A | -1,58 | 0,249 | 5,30 | 0,016 |
| HNF1A | -9,90 | 0,306 | 8,35 | 0,228 |
| ORM1 | -1,48 | 0,420 | 10,55 | 0,009 |
| RBP4 | -2,13 | 0,028 | 3,86 | 0,030 |
| APOE | -1,96 | 0,442 | 4,77 | 0,027 |
| TTR | -1,23 | 0,892 | 5,84 | 0,096 |
| APOC1 | -1,39 | 0,725 | 5,03 | 0,044 |
| GSTA4 | -2,64 | 0,209 | 6,38 | 0,209 |
| CYP1A2 | -2,17 | 0,773 | 2,75 | 0,527 |
| CYP3A4 | 1,06 | 0,979 | -3,18 | 0,384 |
| CEBPA | -2,23 | 0,314 | 2,55 | 0,440 |
| FOXA1 | -11,80 | 0,011 | 3,83 | 0,290 |
| AGR2 | -1,03 | 0,903 | -4,55 | 0,319 |
| CLDN10 | 1,33 | 0,529 | -23,53 | 0,031 |
| CLDN4 | 61,14 | 0,010 | -3,37 | 0,049 |
| MMP7 | 14,28 | 0,301 | -131,42 | 0,013 |
| MUC5B | 1,38 | 0,741 | -3,81 | 0,334 |
| TACSTD2 | 9,89 | 0,018 | -10,54 | 0,003 |
| CFTR | 13,14 | 0,189 | -57,67 | 0,006 |
| PROM1 | -1,83 | 0,646 | -95,73 | 0,025 |
| ST14 | 17,43 | 0,181 | -4,35 | 0,097 |
| KRT19 | 5,22 | 0,437 | -7,71 | 0,255 |
| KRT7 | 7,92 | 0,062 | -67,43 | 0,001 |
| SPP1 | 6,19 | 0,010 | 1,22 | 0,725 |
| SOX9 | 13,68 | 0,128 | 2,35 | 0,495 |
| EPCAM | -14,66 | 0,207 | -42,17 | 0,019 |
| CDH6 | 6,32 | 0,426 | -8,84 | 0,142 |
| CTNND2 | 1,33 | 0,529 | -359,26 | 0,000 |
| NCAM1 | -1,03 | 0,903 | -9,47 | 0,021 |
| SFRP5 | 2,16 | 0,454 | -6,79 | 0,137 |

| **Supplementary Table 6. Pathway’s enrichment between the three microdissected populations** | | | |
| --- | --- | --- | --- |
| **Pathway** | **p HB vs HC** | **p DR vs HC** | **p DR vs HB** |
| KEGG_CITRATE_CYCLE_TCA_CYCLE | 0,049 | 0,004 | 0,010 |
| KEGG_FATTY_ACID_METABOLISM | 0,412 | 0,017 | 0,003 |
| KEGG_PRIMARY_BILE_ACID_BIOSYNTHESIS | 0,422 | 0,035 | 0,018 |
| KEGG_RETINOL_METABOLISM | 0,456 | 0,030 | 0,019 |
| KEGG_DRUG_METABOLISM_OTHER_ENZYMES | 0,447 | 0,032 | 0,010 |
| PID_HNF3B_PATHWAY | 0,118 | 0,003 | 0,007 |
| REACTOME_BILE_ACID_AND_BILE_SALT_METABOLISM | 0,290 | 0,021 | 0,010 |
| REACTOME_GLUCONEOGENESIS | 0,291 | 0,006 | 0,015 |
| HALLMARK_BILE_ACID_METABOLISM | 0,983 | 0,010 | 0,001 |
| BIOCARTA_CASPASE_PATHWAY | 0,044 | 0,502 | 0,589 |
| REACTOME_INTRINSIC_PATHWAY_FOR_APOPTOSIS | 0,028 | 0,021 | 0,262 |
| REACTOME_ACTIVATION_OF_BAD_AND_TRANSLOCATION_TO_MITOCHONDRIA | 0,031 | 0,070 | 0,958 |
| REACTOME_CYTOCHROME_C_MEDIATED_APOPTOTIC_RESPONSE | 0,001 | 0,029 | 0,387 |
| REACTOME_APOPTOTIC_FACTOR_MEDIATED_RESPONSE | 0,038 | 0,041 | 0,994 |
| HALLMARK_APOPTOSIS | 0,021 | 0,820 | 0,681 |
| KEGG_LEUKOCYTE_TRANSENDOTHELIAL_MIGRATION | 0,022 | 0,091 | 0,831 |
| PID_NFKAPPAB_CANONICAL_PATHWAY | 0,043 | 0,087 | 0,781 |
| PID_CD40_PATHWAY | 0,002 | 0,484 | 0,692 |
| PID_TNF_PATHWAY | 0,048 | 0,515 | 0,984 |
| REACTOME_CHEMOKINE_RECEPTORS_BIND_CHEMOKINES | 0,065 | 0,026 | 0,728 |
| REACTOME_INTERFERON_SIGNALING | 0,031 | 0,966 | 0,403 |
| KEGG_JAK_STAT_SIGNALING_PATHWAY | 0,008 | 0,047 | 0,624 |
| KEGG_PATHWAYS_IN_CANCER | 0,006 | 0,191 | 0,387 |
| PID_INTEGRIN3_PATHWAY | 0,006 | 0,058 | 0,508 |
| PID_ILK_PATHWAY | 0,024 | 0,832 | 0,346 |
| PID_CXCR4_PATHWAY | 0,002 | 0,138 | 0,868 |
| PID_INTEGRIN5_PATHWAY | 0,004 | 0,131 | 0,523 |
| REACTOME_EXTRACELLULAR_MATRIX_ORGANIZATION | 0,032 | 0,030 | 0,172 |
| REACTOME_ACTIVATION_OF_MATRIX_METALLOPROTEINASES | 0,285 | 0,018 | 0,340 |
| REACTOME_INTEGRIN_SIGNALING | 0,004 | 0,114 | 0,012 |

| **Supplementary Table 7. Functional analysis of up-regulated genes in hepatobiliary cells vs hepatocytes** | | | |
| --- | --- | --- | --- |
| **MSigDB Hallmark and KEGG pathways** | ***p*** | **GO Pathway** | ***p*** |
| KRAS signaling up | 0.001 | G2/M transition of mitotic cell cycle (GO:0000086) | 0.001 |
| Leukocyte transendothelial migration | 0.001 | cell cycle G2/M phase transition (GO:0044839) | 0.002 |
| TNF-alpha signaling via NF-KB | 0.003 | amide transport (GO:0042886) | 0.002 |
| Toll-like receptor signaling pathway | 0.006 | vitamin transport (GO:0051180) | 0.004 |
| ErbB signaling pathway | 0.008 | mitotic cell cycle phase transition (GO:0044772) | 0.004 |
| Chemokine signaling pathway | 0.018 | Chemokine (C-X-C motif) ligand 12 signaling pathway (GO:0038146) | 0.004 |
| Epithelial mesenchymal transition | 0.022 |  |  |
| Cellular senescence | 0.042 |  |  |
| Inflammatory response | 0.054 |  |  |

| **Supplementary Table 8. Functional analysis of up-regulated genes in hepatobiliary cells vs ductular reaction cells** | | | |
| --- | --- | --- | --- |
| **MSigDB Hallmark and KEGG pathways** | ***p*** | **GO Pathway** | ***p*** |
| Pancreatic secretion | 0.000 | ion homeostasis (GO:0050801) | 0.000 |
| Bile secretion | 0.002 | cellular response to ketone (GO:1901655) | 0.000 |
| Notch signaling pathway | 0.004 | inorganic cation import across plasma membrane (GO:0098659) | 0.001 |
| Apical Junction | 0.016 | cellular response to forskolin (GO:1904322) | 0.001 |
| Wnt-beta Catenin Signaling | 0.023 | response to vitamin D (GO:0033280) | 0.001 |
|  |  | cellular response to alcohol (GO:0097306) | 0.001 |
|  |  | cell chemotaxis (GO:0060326) | 0.002 |
|  |  | regulation of signal transduction (GO:0009966) | 0.002 |
|  |  | regulation of Wnt signaling pathway (GO:0030111) | 0.002 |

| **Supplementary Table 9. Hepatobiliary cells gene signature correlation with clinical parameters** | | | | | |
| --- | --- | --- | --- | --- | --- |
| **GENE** | **Fc HB vs HC** | ***p* SAH vs AH** | **p-CHILD** | **p-MELD** | **ABIC** |
| IER5L | 374,045 | 0,008 |  |  | * |
| MMP2 | 364,15 | 0,025 | * |  | * |
| TGFBR1 | 204,29 | 0,006 | * | * | * |
| UBASH3B | 188,93 | 0,008 | * |  | * |
| CXCR4 | 112,31 | 0,018 | * |  |  |
| PIK3R5 | 73,04 | 0,001 | * | * | * |
| TPM4 | 66,71 | 0,016 | * |  |  |
| CD22 | 56,91 | 0,018 |  |  | * |
| STK39 | 56,23 | 0,000 | * | * | * |
| SFN | 55,87 | 0,004 | * | * |  |
| WBP5 | 54,60 | 0,024 | * |  |  |
| CTHRC1 | 54,46 | 0,007 | * |  | * |
| NCEH1 | 50,42 | 0,007 | * | * | * |
| IGSF3 | 48,17 | 0,015 | * |  |  |
| HES4 | 45,39 | 0,049 | * |  | * |
| SLC12A8 | 44,79 | 0,021 |  |  | * |
| KLF4 | 37,64 | 0,014 | * |  | * |
| ANTXR1 | 37,10 | 0,038 | * |  | * |
| MOXD1 | 32,18 | 0,025 | * |  | * |
| SLC2A6 | 27,51 | 0,000 |  | * | * |
| SRGAP1 | 25,89 | 0,004 | * |  | * |
| LAIR1 | 22,49 | 0,002 | * |  | * |
| ADRBK2 | 16,00 | 0,001 | * | * | * |
| TC2N | 14,91 | 0,041 | * |  | * |
| CCDC109B | 11,98 | 0,019 | * |  | * |
| SH3PXD2B | 8,69 | 0,001 | * |  |  |
| C12orf75 | 7,83 | 0,032 | * |  |  |
| CXCL8 | 7,30 | 0,008 | * | * | * |
| S100A11 | 6,48 | 0,001 | * |  | * |
| SPP1 | 6,19 | 0,037 | * |  | * |
| CHST11 | 5,13 | 6,54E-06 | * | * | * |
| LCN2 | 4,39 | 0,006 | * |  | * |
| RBM11 | 3,88 | 0,011 | * | * | * |
| ERC2 | 3,88 | 0,003 | * | * | * |
| CXCL1 | 3,59 | 0,005 | * | * |  |
| PMP22 | 3,29 | 0,005 | * | * | * |
| LIMK2 | 3,29 | 0,001 | * | * | * |
| FMNL2 | 3,21 | 0,000 | * | * | * |
| RELB | 2,27 | 0,002 |  | * | * |

| **Table S10. Gene expression profile of CXCR4 pathway genes in hepatobiliary cells Vs. hepatocytes** | |
| --- | --- |
| **GENE** | **Fc HB vs HC** |
| **CXCR4** | 112,31 |
| **PIK3R5** | 73,04 |
| **PAG1** | 71,78 |
| **VAV1** | 50,87 |
| **ITGA4** | 49,29 |
| **CSK** | 48,98 |
| **ITGA8** | 46,22 |
| **SSH1** | 41,31 |
| **RGS1** | 29,58 |
| **PTPN11** | 23,50 |
| **PTPRC** | 20,34 |
| **HCK** | 15,30 |
| **GNG2** | 14,97 |
| **HLA-DRB1** | 14,56 |
| **PIK3CD** | 13,75 |
| **PLCB2** | 11,89 |
| **JAK2** | 11,40 |
| **GNAO1** | 9,52 |
| **FGR** | 9,41 |
| **ARL17B** | 7,73 |
| **MLST8** | 7,60 |
| **RALB** | 7,23 |
| **CXCL12** | 5,90 |
| **UBE2N** | 5,71 |
| **PIK3CG** | 5,48 |
| **PLCB1** | 5,02 |
| **LCK** | 4,35 |
| **STAT5A** | 3,99 |
| **ITCH** | 3,92 |
| **PRR4** | 3,64 |
| **LIMK1** | 3,61 |
| **PIK3R6** | 3,61 |
| **PTPN6** | 3,16 |

HB, hepatobiliary cell; HC, hepatocyte

[1]	Aguilar‐Bravo B, Rodrigo‐Torres D, Ariño S, Coll M, Pose E, Blaya D, et al. Ductular Reaction Cells Display an Inflammatory Profile and Recruit Neutrophils in Alcoholic Hepatitis. Hepatology 2019;69:2180–95. https://doi.org/10.1002/hep.30472.

[2]	Aguilar-Bravo B, Sancho-Bru P. Laser capture microdissection: techniques and applications in liver diseases. Hepatol Int 2019;13:138–47. https://doi.org/10.1007/s12072-018-9917-3.

[3]	López-Vicario C, Alcaraz-Quiles J, García-Alonso V, Rius B, Hwang SH, Titos E, et al. Inhibition of soluble epoxide hydrolase modulates inflammation and autophagy in obese adipose tissue and liver: role for omega-3 epoxides. Proc Natl Acad Sci U S A 2015;112:536–41. https://doi.org/10.1073/PNAS.1422590112.

[4]	Hu H, Gehart H, Artegiani B, LÖpez-Iglesias C, Dekkers F, Basak O, et al. Long-Term Expansion of Functional Mouse and Human Hepatocytes as 3D Organoids. Cell 2018;175:1591-1606.e19. https://doi.org/10.1016/j.cell.2018.11.013.

[5]	Argemi J, Latasa MU, Atkinson SR, Blokhin IO, Massey V, Gue JP, et al. Defective HNF4alpha-dependent gene expression as a driver of hepatocellular failure in alcoholic hepatitis. Nat Commun 2019. https://doi.org/10.1038/s41467-019-11004-3.

[6]	Rodrigo-Torres D, Affò S, Coll M, Morales-Ibanez O, Millán C, Blaya D, et al. The biliary epithelium gives rise to liver progenitor cells. Hepatology 2014;60:1367–77. https://doi.org/10.1002/hep.27078.
